# Sticks and Stones, a conserved cell surface ligand for the Type IIa RPTP Lar, regulates neural circuit wiring in *Drosophila*

**DOI:** 10.1101/2020.11.03.367540

**Authors:** Namrata Bali, Hyung-Kook (Peter) Lee, Kai Zinn

**Author notes:** Corresponding author: Kai Zinn, Namrata Bali.

## Abstract

Control of tyrosine phosphorylation is an essential element of many cellular processes, including proliferation, differentiation neurite outgrowth, and synaptogenesis. Receptor-like protein-tyrosine phosphatases (RPTPs) have cytoplasmic phosphatase domains and cell adhesion molecule (CAM)-like extracellular domains that interact with cell-surface ligands and/or co-receptors. We identified a new ligand for the *Drosophila* Lar RPTP, the immunoglobulin superfamily CAM Sticks and Stones (Sns). Lar is orthologous to the three Type IIa mammalian RPTPs, PTPRF (LAR), PTPRD (PTPδ), and PTPRS (PTPσ). Lar and Sns bind to each other in embryos and *in vitro*. The human Sns ortholog, Nephrin, binds to PTPRD and PTPRF. Genetic interaction studies show that Sns is essential to Lar’s functions in several developmental contexts in the larval and adult nervous systems. In the larval neuromuscular system, *Lar* and *sns* transheterozygotes (*Lar/sns* transhets) have synaptic defects like those seen in *Lar* mutants and Sns knockdown animals. Lar and Sns reporters are both expressed in motor neurons and not in muscles, so Lar and Sns likely act in *cis* (in the same neurons). *Lar* mutants and *Lar*/*sns* transhets have identical axon guidance defects in the larval mushroom body in which Kenyon cell axons fail to stop at the midline and do not branch. Pupal Kenyon cell axon guidance is similarly affected, resulting in adult mushroom body defects. Lar is expressed in larval and pupal Kenyon cells, but Sns is not, so Lar-Sns interactions in this system must be in *trans* (between neurons). Lastly, R7 photoreceptor axons in *Lar* mutants and *Lar/sns* transhets fail to innervate the correct M6 layer of the medulla in the optic lobe. Lar acts cell-autonomously in R7s, while Sns is only in lamina and medulla neurons that arborize near the R7 target layer. Therefore, the Lar-Sns interactions that control R7 targeting also occur in *trans*.

## Introduction

Neural circuit assembly is a complex process that involves axon pathfinding, target selection, and synaptogenesis with appropriate synaptic targets. Many cell-surface and secreted proteins are involved in one or more of these stages. Cell adhesion molecules (CAMs) are one class of molecules that play important roles in the process of circuit assembly. Many CAMs are localized to nascent synapses, where they initiate cell-cell contact and recruit pre- and postsynaptic proteins to direct synapse assembly. CAMs typically consist of three domains: an extracellular domain (ECD) that interacts with other CAMs, either homophilically or heterophilically, a transmembrane domain, and an intracellular domain that can interact with cytoplasmic or membrane-associated proteins to regulate signaling. Disruption of CAM activities can lead to neurodevelopmental or neurodegenerative diseases such as autism and schizophrenia (Sudhof, 2008). Some well-known CAMs involved in circuit assembly include N-Cadherin (Bulgakova et al., 2012; Prakash et al., 2005), Dscam (Hattori et al., 2007; Hattori et al., 2008), Neurexins, and Neuroligins (Sudhof, 2008).

Receptor-like protein tyrosine phosphatases (RPTPs) are signaling receptors with CAM-like ECDs that bind heterophilically to ligands, and cytoplasmic domains with tyrosine phosphatase enzymatic activity (reviewed by (Coles et al., 2015)). Type IIa (R2A), IIb (R2B) and III (R3) RPTP subtypes have ECDs containing immunoglobulin superfamily (IgSF) domains and/or fibronectin type III (FNIII) repeats. *Drosophila* has six RPTPs, four of which are expressed primarily in the nervous system. Lar is a Type IIa RPTP with both IgSF and FNIII domains that is orthologous to a 3-member family of mammalian RPTPs: PTPRF (Lar), PTPRD (PTPδ, R-PTP-δ), and PTPRS (PTPσ, R-PTP-σ). *C. elegans* has a single Lar ortholog, PTP-3. Ptp69D resembles Type IIa RPTPs, having IgSF and FNIII domains, but does not correspond to any specific mammalian RPTP. Ptp10D and Ptp99A have only FNIII repeats in their ECDs. There are also two other RPTPs: Ptp52F, which is primarily expressed in the gut but also has functions in neurons, and Ptp4E, which is ubiquitously expressed. Ptp10D and Ptp4E are orthologous to mammalian Type III RPTPs, which are regulators of receptor tyrosine kinases (reviewed by (Jeon and Zinn, 2015)). There are no Type IIb RPTPs in *Drosophila*.

Drosophila Lar and Ptp69D have well-characterized functions during neural development. Motor axon guidance in embryos is altered in mutants for both RPTPs. They genetically interact with each other and with the other four RPTPs. In double, triple, and quadruple mutants there are specific alterations in motor and central nervous system (CNS) axon guidance (Jeon et al., 2008; Sun et al., 2000). *Lar* mutations affect the structures of neuromuscular junctions (NMJs) in the larval neuromuscular system (Johnson et al., 2006; Kaufmann et al., 2002). Lar is also required for development of the larval mushroom body (MB) lobes (Kurusu and Zinn, 2008) and for R7 photoreceptor axon targeting in the optic lobe (OL) (Clandinin et al., 2001; Maurel-Zaffran et al., 2001). Ptp69D works together with Lar to control R7 axon guidance and targeting (Hakeda-Suzuki et al., 2017; Hofmeyer and Treisman, 2009).

Mammalian Type IIa RPTPs function during synaptogenesis, and knockouts of the three genes have developmental defects causing craniofacial malformations (Stewart et al., 2013), severe muscle dysgenesis and loss of motor neurons in the spinal cord (Uetani et al., 2006). *Ptprs* and *Ptprd* mutant mice exhibit increased paired pulse facilitation, enhanced or reduced long-term potentiation, respectively, and distinct behavioral alterations (Horn et al., 2012; Uetani et al., 2000). *Ptprs* and *Ptprd* mutant mice also exhibit early growth retardation and increased neonatal mortality (Uetani et al., 2000). The chicken orthologs of PTPRD and PTPRS are expressed in developing motor neurons, and knockdown of each of these PTPs affected the dorsal nerve, causing either abnormal fasciculation or a reduced or missing nerve (Stepanek et al., 2005).

The CAM-like structure of the Lar ECD implies that its functions are regulated by interactions with cell-surface or extracellular ligands. We and others identified the heparan sulfate proteoglycans (HSPGs) Syndecan (Sdc) and Dally-like (Dlp) as Lar ligands (Fox and Zinn, 2005; Johnson et al., 2006). Lar interacts directly with heparan sulfate, as do mammalian Type IIa RPTPs (Aricescu et al., 2002). Sdc and Dlp have genetic interactions with Lar (Fox and Zinn, 2005; Johnson et al., 2006). They are involved in Lar’s regulation of embryonic axon guidance and larval NMJ development, but *Sdc* and *Dlp* phenotypes are much weaker than *Lar* phenotypes, implying that other ligands are also involved. The HSPGs are not involved in regulation of Lar’s functions in R7 photoreceptor axon guidance (Hofmeyer and Treisman, 2009).

A number of protein ligands for mammalian Type IIa RPTPs have been identified. These include Netrin-G ligand 3 (NGL-3) (Kwon et al., 2010; Woo et al., 2009), Tropomyosin kinase C (TrkC) (Takahashi et al., 2011), Interleukin-1 receptor accessory protein-like 1 (IL1RAPL1) (Yoshida et al., 2011), Interleukin-1 receptor accessory protein (IL-1RAcP) (Yoshida et al., 2012), Slit- and Trk-like family protein (Slitrk) 1-Slitrk6 (Takahashi et al., 2012; Yim et al., 2013), synaptic adhesion-like molecule (SALM) 3, and SALM5 (Choi et al., 2016; Li et al., 2015; Wojtowicz et al., 2020). Each of these ligands localizes to postsynaptic membranes, where they form heterophilic *trans* complexes with presynaptic RPTPs to regulate cell-cell adhesion, presynaptic differentiation and excitatory synapse development (Takahashi and Craig, 2013).

Here we describe the identification of Sticks and Stones (Sns) as a new Lar ligand, and show that the Lar-Sns interaction is conserved between flies and mammals. Sns is a single-pass transmembrane protein with a large ECD with IgSF and FNIII domains. Sns has orthologs in *C. elegans* (SYG-2) and mammals (Nephrin). It belongs to a subfamily of IgSF proteins called Irre cell recognition module (IRM) proteins, which has four members in *Drosophila*: Sns, Kirre, Roughest (Rst), and Hibris (Hbs) (Fischbach et al., 2009). Kirre and Rst are paralogs that bind to Sns and Hbs, as well as to each other (Bour et al., 2000; Galletta et al., 2004; Ozkan et al., 2013; Shelton et al., 2009). The *C. elegans* orthologs of Sns and Kirre, SYG-2 and SYG-1, also bind to each other (Ozkan et al., 2014). In humans and mice, there is one Sns/Hbs ortholog, Nephrin, and three Kirre/Rst orthologs (Kirrels or Nephs).

The Sns ECD contains nine Ig domains and a single FNIII repeat, while the Kirre and Rst ECDs contain five Ig domains (Ozkan et al., 2014). All four IRM proteins function together as ligand-receptor pairs on the surface of founder cells and fusion competent myoblasts to regulate myoblast fusion (Bour et al., 2000; Shelton et al., 2009). The four proteins also function together in nephrocyte development (Zhuang et al., 2009) and in ommatidium patterning in the retina (Bao et al., 2010). *C. elegans* SYG-2 and SYG-1 regulate the formation of synapses by HSNL neuron onto vulval muscles (Ozkan et al., 2014; Shen, 2004). SYG-1 acts presynaptically, while SYG-2 acts in the guidepost epithelial cells to direct presynaptic component assembly at the site of their interaction (Shen, 2004). The mammalian orthologs of Sns and Kirre, Nephrin and Neph1/Kirrel1, are required for the formation and functioning of the kidney slit diaphragm (Liu et al., 2003; Ruotsalainen et al., 1999).

In this paper, we show that interactions with Sns control Lar’s functions in MB lobe development and R7 photoreceptor axon targeting. Transheterozygous animals lacking one copy of the wild-type *Lar* and *sns* genes have MB and R7 phenotypes identical to those of *Lar* homozygotes. The relevant Lar-Sns interactions appear to be in *trans*, because Lar and Sns are expressed in different neurons. Sns also regulates Lar’s functions in NMJ development, but in that system the two proteins are expressed in the same neurons and likely interact in *cis*.

## Results

### Identification of Sns as a Lar binding partner

Cell-surface protein (CSP) interactions mediated by ECDs are often of low affinity, having K_d_s in the micromolar range and fast dissociation rates. This usually precludes identification of such interactions through methods such as affinity purification/mass spectrometry (AP/MS), since they do not form stable complexes. Low-affinity CSP-CSP interactions create stable adhesive interactions between cells through avidity effects, since there are many copies of each protein at cell interfaces. Successful *in vitro* detection of low-affinity CSP interactions often requires taking advantage of avidity through clustering. Clustering is achieved by incorporating multimerization domains into soluble ECD fusion proteins, which can be used for ELISA-based assays and cell staining. The RPTPs were included in an ELISA-based global extracellular interactome (ECIA) screen of all *Drosophila* cell-surface proteins containing IgSF and/or FNIII domains (Ozkan et al., 2013). However, interactions above background were not detected for any RPTP in this screen, which was conducted using unpurified cell supernatants.

We developed the live-dissected embryo staining screen as a way to identify low-affinity binding partners among neural CSPs in an assay in which they are expressed in a normal cellular context. This screen may be more sensitive than the ECIA because it takes maximum advantage of avidity effects. Multimeric ECD fusion proteins are added to live-dissected embryos and incubated for ~2 hours at room temperature, during which time complexes of fusion proteins with overexpressed CSPs can coalesce (“cap”) into dense patches. The embryos are then washed directly with paraformaldehyde, which crosslinks these patches and freezes complexes into place. The complexes are then visualized with fluorescent secondary antibody (Fox and Zinn, 2005; Lee et al., 2009).

We first used this method to identify the heparan sulfate proteoglycan (HSPG) Syndecan (Sdc) as a ligand for Lar, using a deficiency (*Df*) screen to find a region of the genome whose deletion eliminated Lar ECD staining and then narrowing down the region to a single gene, *Sdc* (Fox and Zinn, 2005). We later developed a gain-of-function (GOF) version of the screen in which we crossed 300 lines bearing “EP-like” (UAS-containing) *P* elements upstream of CSP genes to a strong pancellular driver, tubulin (tub)-GAL4. This collection of lines had first been used for an *in vivo* screen in which we crossed each line to a muscle GAL4 driver and then searched for motor axon phenotypes conferred by high-level muscle expression (Kurusu et al., 2008). Using the GOF screen, we identified Stranded at second (Sas), a large CSP expressed in epidermal cells, as a ligand for Ptp10D (Lee et al., 2013).

To conduct the GOF screen with Lar ECD fusion proteins, we first needed to eliminate Lar binding to Sdc, which is mediated by short basic sequences in the first Lar Ig domain that bind to heparan sulfate (HS). Accordingly, for the screen described here, we used a mutant, HS2, that eliminates HS binding (Fox and Zinn, 2005). We expressed a dimeric LAR^HS2^-alkaline phosphatase (henceforth called Lar-AP) fusion protein at high levels using the baculovirus system and stained live-dissected stage 16 embryos with unpurified supernatant, which reduces background. We screened crosses for all 300 EP-like lines to tub-GAL4 in this manner.

As shown in Figs. 1A-B, we observed very faint CNS axon staining in wild-type (WT) embryos, but saw bright staining in both the CNS axon ladder and the periphery when tub-GAL4 was crossed to a line that has an insertion of an EP-like element ~200 bp 5’ to the transcription start of *sns (sns^EY08142^)*. This indicates that ectopic expression of Sns in neural, ectodermal, and muscle cells confers binding to the Lar ECD. However, it does not prove that Lar and Sns bind directly to each other, since the results could also be explained if Sns ectopic expression induced expression or stabilization of another protein that actually binds to Lar.

**Figure 1:**
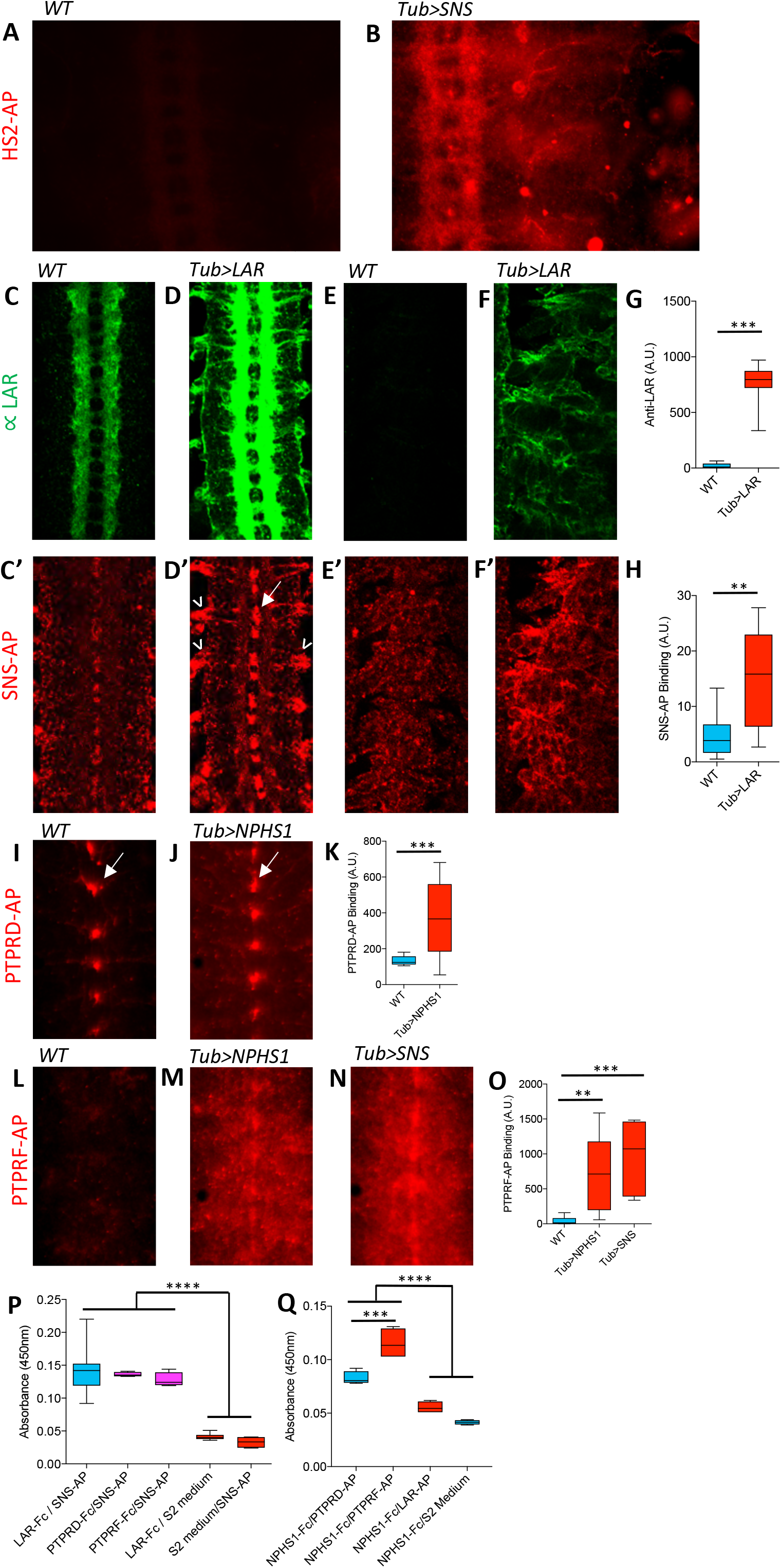
Binding of Lar and its orthologs to Sns and Nephrin. All images show live-dissected late stage 16 embryos. (A, B) Staining with HS2-AP, a heparan-sulfate binding deficient version of LAR-AP (Fox and Zinn), visualized with anti-AP antibody. (A) WT embryo; HS2-AP weakly binds to CNS axons. (B) Tub>Sns embryo, showing bright ectopic staining by HS2-AP in the CNS and periphery. (C-F) Lar overexpression in Tub>Lar embryos, visualized with anti-Lar mAb. (C, D) CNS axon staining in WT (C) and Tub>Lar (D). Longitudinal axons are stained in WT; all axons are brightly stained in Tub>Lar. (E, F) Staining in the periphery in WT (E) and Tub>Lar (F). There is no visible staining in WT (motor axon staining is below the level of detection), while Tub>Lar embryos show widespread staining. (G) Quantitation of CNS staining with anti-Lar in WT and Tub>Lar. (C’-F’) Staining with Sns-AP_5_ in WT and Tub>Lar embryos, visualized with anti-AP antibody. (C’, D’) CNS staining in WT (C’) and Tub>Lar (D’). Midline glia and cell bodies are weakly stained in WT (C’); note that this pattern does not resemble anti-Lar staining (C). Midline glia and exit junctions are brightly stained in Tub>Lar (D’); note the similarity between the exit junction patterns visualized with anti-Lar (D) and Sns-AP. (E’, F’) Staining in the periphery in WT (E’) and Tub>Lar (F’). Widespread staining in the periphery is increased in intensity in Tub>Lar. (H) Quantitation of CNS staining with Sns-AP in WT and Tub>Lar. (I, J) CNS staining with PTPRD-AP_5_ in WT (I) and Tub>NPHS1 (J). Note midline glial staining in WT; this staining is not increased in intensity in Tub>NPHS1, but staining intensity in the remainder of the CNS is increased. (K) Quantitation of CNS staining in WT and Tub>NPHS1. (L-N) CNS staining with PTPRF-AP_5_ in WT (L), Tub>NPHS1 (M), and Tub>Sns (N) embryos. Note that there is very little staining in WT, but bright staining in the entire CNS in Tub>NHS1 and Tub>Sns. (O) Quantitation of CNS staining in WT, Tub>NPHS1, and Tub>Sns. (P, Q) *In vitro* binding measured by modified ELISA assay, using HRP enzymatic activity for detection. (P) Sns-AP_5_ prey binds to Lar-Fc, PTPRD-Fc, and PTPRF-Fc baits, Sns-AP_5_ prey does not bind to S2 medium bait, and there is no signal with Lar-Fc bait and S2 medium prey. (Q) PTPRF-AP_5_ prey bind to Nephrin-Fc (NPHS1-Fc) bait, but Lar-AP_5_ does not. There is no signal with Nephrin-Fc bait and S2 medium prey. There is also a small increase in signal over background with Nephrin-Fc bait and PTPRD-AP_5_ prey.

To address this issue, we performed “reverse-binding” experiments to determine whether Lar binds directly to Sns. To do this, we used tub-GAL4 to drive pancellular expression of Lar in embryos, using a transgenic UAS-Lar construct. To confirm that Lar was overexpressed, we stained WT and tub>Lar stage 16 embryos with anti-Lar antibody. In Tub>Lar embryos, Lar was strongly expressed on CNS axons as well as in the periphery, while in WT embryos Lar is localized to longitudinal axon tracts and there is no expression in the periphery (Figs. 1C-F).

We then stained WT and tub>Lar embryos with a pentameric Sns-AP fusion protein, containing the ECD of Sns fused to a COMP pentamerization domain and AP (Ozkan et al., 2013). Sns-AP_5_ stained the CNS in WT embryos (Fig. 1C’). Interestingly, midline glia (arrow) were more strongly stained than axons and cell bodies. This pattern does not resemble Lar antibody staining (Fig. 1C), indicating that Sns has another binding partner in embryos, perhaps Kirre or Rst. There is also weak Sns-AP_5_ staining in the periphery (Fig. 1E’), where Lar is not expressed.

Embryos with ectopic expression of Lar driven by tub-GAL4 showed a three-fold increase in Sns-AP_5_ staining in the CNS compared to WT control embryos (Figs. 1D’, H). Sns-AP_5_ staining was increased at sites where motor axons exit the ventral nerve cord (VNC; arrowhead, Figure 1D’) and in midline glia. Staining was also increased in the periphery (Figs. 1E’, F’). Some of this ectopic staining colocalizes with ectopic Lar, such as at the VNC exit points. However, there is only weak staining on CNS axons, where Lar is most prominently expressed. This indicates that either Lar on axons is inaccessible to Sns-AP_5_, or that there is another protein expressed in midline glia and exit junctions that facilitates Sns-AP_5_ binding to Lar. In any case, the reverse binding experiment provides strong evidence that Lar and Sns directly interact with each other.

To confirm direct binding, we conducted ECIA experiments with dimeric Lar-Fc and pentameric Sns-AP_5_ proteins made in Schneider 2 (S2) cells. Sns-AP_5_ “prey” exhibited specific binding to Lar-Fc “bait” coupled to the surface of an ELISA plate, showing that the two proteins do interact directly *in vitro* (Fig. 1P). This binding signal is weak compared to what is typically observed for strongly interacting partners such as Sns and Kirre, being only 3-fold over background, but is nevertheless statistically significant. This may indicate that Lar-Sns interactions are of low affinity (tens or hundreds of micromolar).

### Mammalian orthologs of Lar and Sns bind to each other in embryos and *in vitro*

Lar is orthologous to the Type IIa RPTP subfamily, which has three members in mouse and human: PTPRF, PTPRD, and PTPRS. Sns is orthologous to Nephrin. To determine whether binding between Lar and Sns is evolutionarily conserved, we tested whether Nephrin binds to PTPRD, PTPRS or PTPRF in live-dissected embryos. We made a transgenic line with a UAS-linked full-length human Nephrin cDNA (NPHS1) construct. We made AP_5_ fusion proteins containing the ECDs of PTPRD, PTPRS and PTPRF in S2 cells.

We tested the binding patterns of each of the three AP fusion proteins in WT, tub>NPHS1, and tub>Sns embryos. PTPRD-AP_5_ gave the clearest signal, with strong staining in midline glia and weak staining in the rest of the VNC (Fig. 1I). When Nephrin was ectopically expressed using tub-GAL4 (tub>NPHS1), midline glial staining was not changed much, but there was a 3-fold increase in staining intensity relative to WT in the VNC as a whole (Figs. 1J, K). These data suggest that a PTPRD binding partner is expressed in WT midline glia, or that midline glial membranes bind nonspecifically to this probe.

PTPRF-AP_5_ showed almost no staining of WT embryos, but stained the VNC and midline glia in tub>NPHS1 and tub>Sns embryos (Figs. 1L-N). Staining intensity was increased by 18-20-fold relative to WT when Nephrin or Sns were expressed (Fig. 1O). PTPRS-AP_5_ showed little staining in WT or tub>NPHS1 embryos.

We then tested PTPRF and PTPRD for binding to Sns and Nephrin *in vitro*. We observed that Fc dimers for both human proteins bound to fly Sns-AP_5_, and the signal-to-noise ratio was about the same as for Lar. PTPRF-AP_5_ also bound to Nephrin-Fc, and there was a small increase over background for Nephrin and PTPRD-AP_5_ (Figs. 1P, Q). In summary, these data indicate that the Lar-Sns interaction is evolutionarily conserved for at least two of the three mammalian Lar orthologs.

### Lar and Sns are co-expressed in larval motor neurons

To characterize the cells that express Lar and Sns, we created T2A-GAL4 lines derived from intronic *MiMIC* insertions in the two genes. We used the method described in (Diao et al., 2015) to insert a T2A-GAL4 “Trojan” exon into *MiMIC* elements in coding introns of *Lar* and *sns*. The expression of Lar and Sns was visualized by crossing the resulting T2A-GAL4 lines to a UAS-EGFP reporter. Note that, in coding intron T2A-GAL4 lines, expression of GAL4 requires in-frame readthrough from the coding region, and therefore reports on the rate of initiation of translation from the correct ATG. Thus, these GAL4s are translational, not just transcriptional, reporters. In the third instar larval VNC, Lar-T2A-GAL4>UAS-EGFP (Lar>GFP) expression was seen in motor neurons (large paired cells) and in a large number of interneurons (Figs. 2A, B). Sns-T2A-GAL4>UAS-EGFP (Sns>GFP) expression was also seen in motor neurons (Figure 2C), as well as in a pattern of interneurons that appeared different from those expressing the Lar reporter (Figure 2D). We stained VNCs for Even-skipped (Eve), which labels the aCC and RP2 motor neurons, and found that the Sns reporter is expressed in RP2 (known as MNISN-1s in larvae; Supp. Fig. 1).

**Figure 2:**
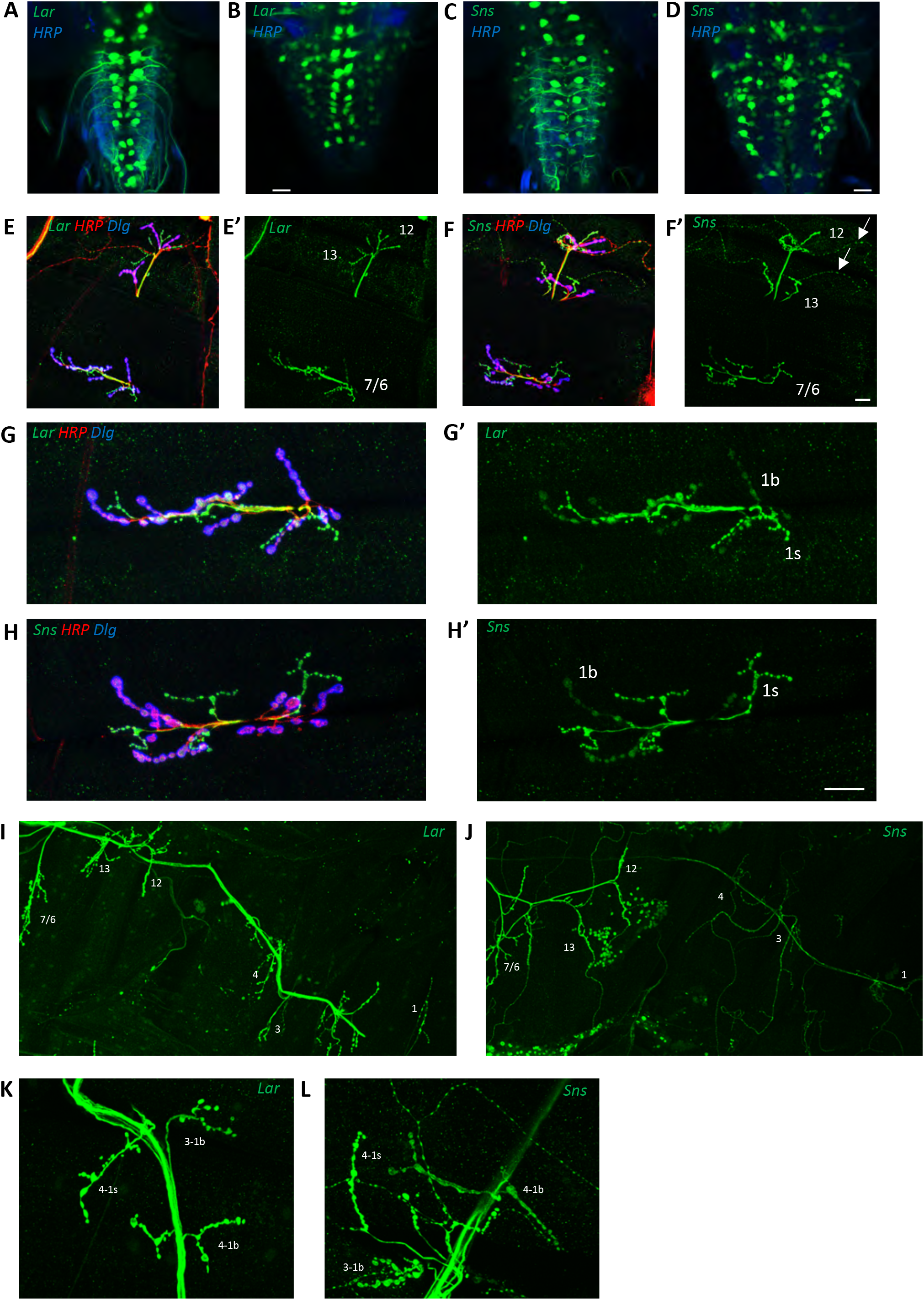
Expression of Lar and Sns reporters in motor neurons. (A-D) Confocal projections of 4-6 optical slices showing EGFP expression driven by either Lar^MI02154-T2A-GAL4^ (Lar>GFP) or Sns^MI03001-T2A-GAL4^ (Sns>GFP) (green) co-stained with anti-HRP (blue). The bright paired midline cells include motor neurons. (E-H’) Confocal projections of larval NMJs on muscles 7/6, 13 and 12 (E-F’) and zoomed-in on muscle 7/6 (G-H’), triple-stained with anti-GFP (green), anti-HRP (red), and anti-Dlg (blue). (E’, F’, G’, H’) show GFP signal only. Anti-HRP labels neuronal membranes, and anti-Dlg labels the subsynaptic reticulum at 1b boutons. Lar>GFP and Sns>GFP expression is seen in both 1b and 1s boutons (green). (I, J) Projection of optical slices through an entire larval hemisegment showing Lar>GFP (I) and Sns>GFP (J) expression in both 1b and 1s motor neurons. Individual muscles are numbered. Dorsal is to the right. Note that Lar>GFP is equally expressed on most axons and NMJs, while Sns>GFP is expressed at lower levels in motor neurons projecting to dorsal muscles. (K, L) Close-up of NMJs on muscles 3 and 4 showing both Lar>GFP (K) and Sns>GFP (L) expression in 1b and 1s NMJs on those muscles. Scale bar, 20 μm.

We characterized Lar and Sns expression in motor neurons by examining larval NMJs labeled with GFP reporter. The 30 body wall muscles in third instar larvae are innervated by 32 motor neurons in each hemisegment. There are two types of glutamatergic motor neurons, 1b and 1s. Each muscle is usually innervated by a single 1b motor neuron. There are two 1s motor neurons, MNISN-1s and MNISNb/d-1s, in each hemisegment. Each 1s motor neuron innervates an entire field of muscles, with MNISN-1s innervating more dorsal muscles including muscles 4, 3, 2 and 1, whereas MNISNb/d-1s innervates more ventral muscles including muscles 7, 6, 13 and 12. In addition to 1b and 1s, two other types of motor neurons innervate a subset of muscles. Each Type II motor neuron innervates several muscles, whereas the Type III motor neuron only innervates muscle 12. We compared the expression of Lar and Sns using their respective T2A-GAL4s in the different types of motor neurons.

Both Lar and Sns were broadly expressed in several different types of motor neurons. Lar>GFP was expressed in most 1b motor neurons, and in both 1s motor neurons (Figs. 2I, K). However, Lar>GFP was not expressed in either Type II or Type III motor neurons (Figs. 2E, E’). Lar>GFP expression was stronger in 1s motor neurons compared to 1b motor neurons. Specifically, at the muscle 7/6 NMJ stronger Lar>GFP expression can be seen in 1s boutons compared to 1b boutons (Fig. 2G, G’). No expression was seen in muscles. These data are consistent with earlier findings on Lar expression (Johnson et al., 2006; Kaufmann et al., 2002).

Sns>GFP was also expressed in both 1b and 1s motor neurons. Expression was stronger in the 1b motor neurons that target more ventral muscles, including muscles 7/6, 13 and 12 (Fig. 2J). Weak Sns>GFP expression could be seen in 1b motor neurons that target more dorsal muscles, including muscles 4 and 3 (Figure 2L). Strong Sns>GFP expression was seen in both 1s motor neurons, MNISN-1s and MNISNb/d-1s (Figure 2J). Similar to Lar>GFP, Sns>GFP was not expressed in the Type III motor neuron. However, unlike Lar>GFP, Sns>GFP expression was seen in Type II motor neurons (Figs. 2F, F’). Similar to Lar>GFP expression being stronger in 1s motor neurons, we observed stronger Sns>GFP expression in 1s boutons compared to the 1b boutons on the same NMJ (Figs. 2H, H’). No Sns>GFP expression was seen in muscles. Thus, Sns is expressed in 1b, 1s and Type II types of motor neurons and is co-expressed with Lar in most 1b and 1s neurons.

### Lar and Sns genetically interact to shape morphogenesis of NMJs

Previous studies have shown that NMJs require appropriate levels of Lar for proper development (Johnson et al., 2006; Kaufmann et al., 2002). Reduction of Lar levels reduces the number of synaptic boutons at the muscle 7/6 NMJ. We asked whether the interaction between Lar and Sns is required for proper development of the NMJ using genetics. Sns is required for myoblast fusion in developing embryos, and *sns* null mutant animals lack body wall muscles and do not survive to the first instar larval stage.

To examine genetic interactions, we combined an *sns* null mutation with two different *Lar* null mutations to analyze NMJ phenotypes in transheterozygote (transhet) animals. *sns*^*xb3*^ is a point mutation resulting in a stop codon in the ECD of the SNS protein. *sns*^*xb3*^ mutant embryos lack Sns protein (Bour et al., 2000). We tested two different alleles of *Lar* with *sns*^*xb3*^: *Lar*^*13.2*^ and *Lar*^*451*^. Both alleles have been described as being presumed null mutations (Clandinin et al., 2001; Krueger et al., 1996). *Lar*^*13.2*^ mutants were shown to display abnormalities at the muscle 7/6 NMJ and in the larval MB (Johnson et al., 2006; Kurusu and Zinn, 2008). *Lar*^*451*^ mutants were characterized for R7 photoreceptor axon guidance defects (Clandinin et al., 2001). We analyzed the NMJs at muscle 7/6 in *Lar*^*13.2*^/*sns*^*xb3*^ and *Lar*^*451*^/*sns*^*xb3*^ transhets and used a semi-automated macro in FIJI to quantify several different parameters at the 7/6 NMJ, including total NMJ area, total NMJ length, longest branch length, number of boutons and number of branches (Nijhof et al., 2016). To dissect out the roles of Lar and Sns in 1b vs. 1s NMJ formation, we performed separate analyses for the 1b and the 1s NMJ arbor at each NMJ.

*Lar*/+ and *sns*/+ heterozygote control animals did not display any overt abnormalities at the 7/6 NMJ (Fig. 3A, B, C). Quantification of 1b NMJ parameters showed no differences between WT and *Lar*^*13.2*^/+, *Lar*^*451*^/+, or *sns*^*xb3*^/+ animals in 1b NMJ area (Fig. 3G), number of 1b boutons (Figure 3H), 1b NMJ length (Figure 3I), longest 1b branch length (Figure 3J) or the number of 1b branches (Figure 3K). On the other hand, the two different *Lar* and *sns* transhets displayed severe NMJ abnormalities, similar to *Lar* null animals (*Lar*^*13.2*^/*Lar*^*451*^). Quantification of NMJ parameters showed that *Lar*^*13.2*^/*sns*^*xb3*^ transhet NMJs had a 34% reduction in 1b NMJ area, 39% reduction in number of 1b boutons, 42% reduced total 1b NMJ length, 33% reduced longest 1b branch length and 45% reduced number of 1b branches (Fig. 3). *Lar*^*451*^/*sns*^*xb3*^ transhets showed slightly stronger phenotypes at the 7/6 NMJ with 48% reduced 1b NMJ area, 64% reduced number of 1b boutons, 57% reduced total 1b NMJ length, 48% reduced longest 1b branch length and 64% reduced number of 1b branches (Fig. 3). Similar reductions in NMJ parameters were observed in *Lar* mutants, which showed 54% reduced NMJ area, 70% reduced number of 1b boutons, 62% reduced total 1b NMJ length, 52% reduced 1b longest branch length, and 72% reduced number of 1b branches. There was no significant difference between the stronger *Lar*^*451*^/*sns*^*xb3*^ transhet and *Lar* mutants for any of the NMJ parameters measured, indicating that Lar and Sns probably function in the same genetic pathway. We also observed similar NMJ abnormalities on other muscles as well, including the muscle 13 and 12 NMJs (data not shown). There was no difference in the size of muscles in the transhets or the *Lar* mutants. This suggests that the Lar-Sns interaction is not required for the role of Sns in myoblast fusion during embryonic development.

**Figure 3:**
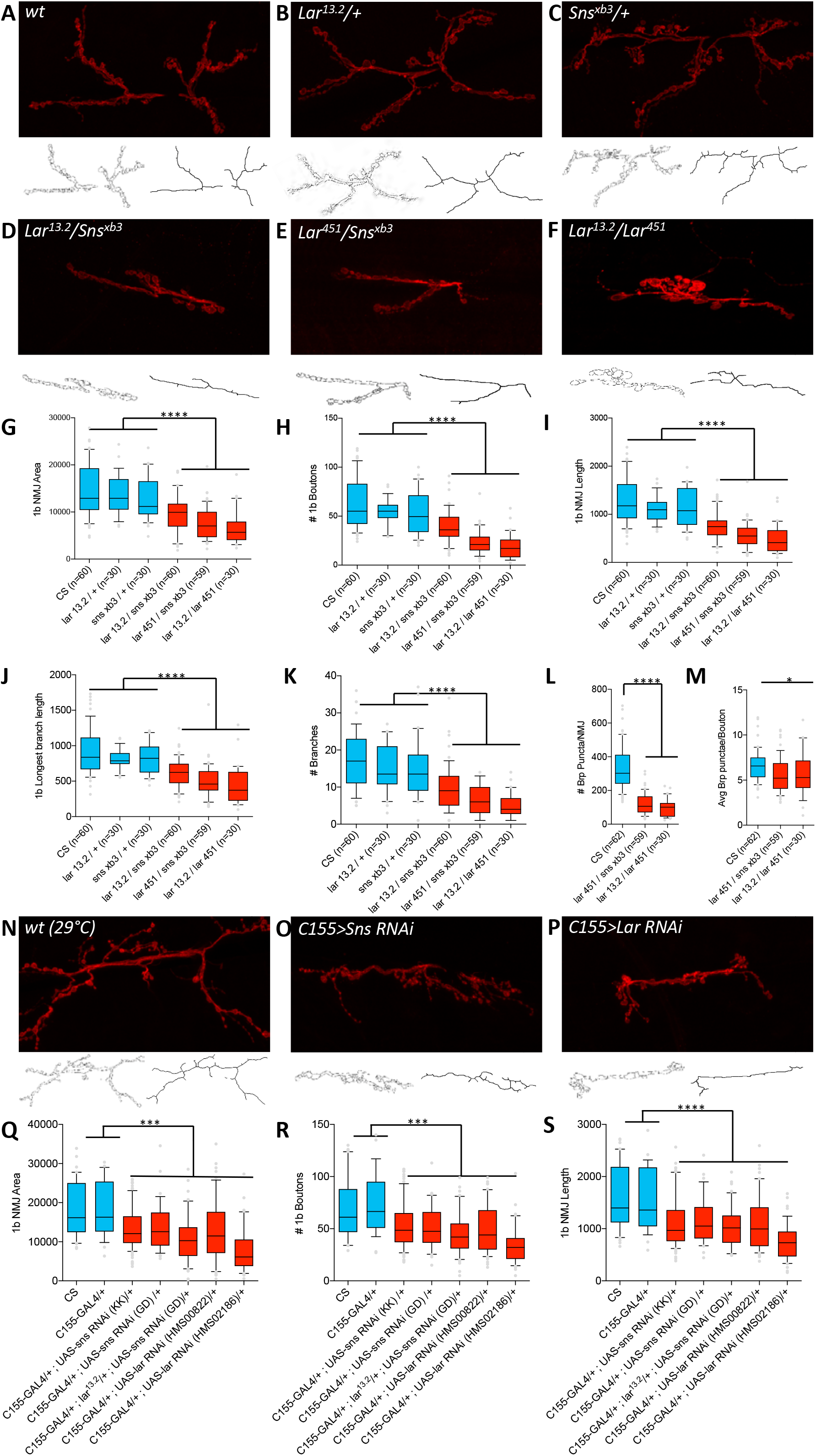
*Lar/sns* transheterozygotes have the same phenotypes as *Lar* mutants and Sns knockdowns. NMJs were analyzed using a published FIJI Macro (Nijhof et al., 2016) that uses HRP to outline boutons and measures NMJ area, perimeter, length, longest branch length, number of branches, number of boutons and Bruchpilot (Brp) labeled punctae. (A-F) Representative images of the NMJ on muscles 7/6 from WT and heterozygote controls (A-C), Lar/sns transheterozygotes (D-E), and *Lar* mutants (F). NMJs are labeled with anti-HRP (red). NMJ outlines showing boutons and branch architecture as outputs from the Macro are under each NMJ image. (G-K) Quantification of 1b NMJ parameters, showing reduced NMJ size and arborization in *Lar/sns* transhets and *Lar* mutants (red) compared to het controls (blue). Data is average from segments A2-A4 from minimum 30 NMJs per genotype. (L, M) Quantification of Brp punctae showing reduced number of active zones in *Lar/sns* transhets and *Lar* mutants. (N-P) Representative images of NMJs on muscles 7/6 from animals with RNAi mediated neuronal knockdown of Lar and Sns. Neuronal *Lar* or *sns* RNAi results in the same NMJ abnormalities seen in genetic *Lar/sns* transhets and *Lar* mutants. (Q-S) Quantification of NMJ parameters showing reduced 1b NMJ area, number of boutons and NMJ length upon either Lar or Sns knockdown. A2-A4 segments were analyzed from at least 30 NMJs on muscles 7/6. All datasets were analyzed using One-Way ANOVA followed by Tukey’s post-hoc correction. **** p<0.0001; *** p<0.001.

We next asked whether the number of synapses was altered in *Lar/sns* transhets and *Lar* mutants at the 7/6 1b NMJ. We used antibodies against the active-zone protein Bruchpilot (Brp) to label active zones in boutons and performed quantitative analyses of Brp-positive punctae using the NMJ FIJI Macro. We quantified the total number of Brp punctae in 1b boutons at the muscle 7/6 NMJ and found that *Lar*^*451*^/*sns*^*xb3*^ transhets and *Lar*^*13.2*^/*Lar*^*451*^ mutants had 64% and 71% fewer Brp punctae per NMJ, respectively, compared to WT NMJs (Fig. 3L). This indicates that there is no compensatory increase in the number of synaptic active zones in response to a reduced NMJ size and number of boutons. During the course of NMJ development, synaptic maturation occurs later than NMJ expansion and arborization. Our results suggest that Lar and Sns are required for both processes of NMJ morphogenesis: initial NMJ expansion and arborization as well as synaptic active zone formation.

Lar and Sns are also required for the morphogenesis of 1s motor neuron arbors at the muscle 7/6 NMJ. 1s NMJs were severely affected in *Lar/sns* transhets and *Lar* mutants. *Lar*^*13.2*^/*sns*^*xb3*^ transhets displayed 50% reduction in 1s NMJ area, 52% reduction in the number of 1s boutons and 46% reduction in the 1s NMJ length (Supp. Fig. 2). *Lar*^*451*^/*sns*^*xb3*^ transhets displayed even stronger phenotypes, with 75% reduced 1s NMJ area, 78% reduced 1s bouton number, and 65% reduced 1s NMJ length (Supp. Fig. 2). *Lar* mutants had similar reductions in 1s NMJ size (65% reduced 1s NMJ area, 82% reduced 1s number of boutons and 82% reduced 1s NMJ length).

We confirmed the NMJ abnormalities seen in *Lar/ sns^xb3^* transhets by analyzing a *sns* deficiency (*Df*) allele which lacks the entire *Sns* gene. We observed similar NMJ abnormalities in *Lar^13.2^/sns^Df^* animals to those seen in *Lar*^*13.2*^/*sns*^*xb3*^and *Lar*^*451*^/*sns*^*xb3*^ transhets (data not shown). Overall, our data shows that interaction between Lar and Sns is required for 1b and 1s NMJ morphogenesis.

### Lar and Sns act in *cis* at the NMJ

While *sns* is expressed in body wall muscles during the period of muscle fusion, its RNA levels decrease in late embryos (Bour et al., 2000). We did not observe any expression of either Lar or Sns reporters in muscles at the third instar larval stage. To confirm that both Lar and Sns act in motor neurons to regulate NMJ development, we performed neuron-specific RNAi knockdown for both Lar and Sns and measured the same NMJ parameters as in the transhets and mutant analyses. We used a pan-neuronal driver, elav^C155^-GAL4 (C155-GAL4) to drive UAS-RNAi lines for either *Lar* or *sns*. We tested two different RNAi lines for both *Lar* and *sns*. Neuronal knockdown of Sns caused NMJ abnormalities similar to those seen in *Lar/sns* transhets and *Lar* mutants (Fig. 3O). Quantification of NMJ parameters revealed that 1b NMJ area was reduced by 25% (*sns* KK RNAi line) and 23% (*sns* GD RNAi line) (Fig. 3Q), the number of 1b boutons was reduced by 22% and 25% in the two *sns* RNAi lines (KK and GD respectively) (Fig. 3R), and 1b total NMJ length was reduced by 28% and 27% in the two RNAi lines, respectively (Fig. 3S).

Neuronal knockdown of Lar caused similar NMJ abnormalities as seen in *Lar/sns* transhets and *Lar* mutants (Fig. 3P). One of the two *Lar* RNAi lines, TRiP HMS00822, showed similar reductions in 1b NMJ area, number of boutons and total NMJ length to those seen with both *sns* RNAi lines, with 1b NMJ area reduced 30%, number of 1b boutons reduced 29% and total NMJ length reduced 30% (Figs. 3Q-S). The second *Lar* RNAi line, TRiP HMS02186, had stronger RNAi effects, as adult flies with neuronal knockdown of Lar mediated by that line were not viable and died during mid-pupal stages. We found stronger effects of Lar knock-down by this RNAi line on NMJ development, with 1b NMJ area reduced 58% (Fig. 3Q), number of 1b boutons reduced 50% (Fig. 3R), and total NMJ length reduced 54% (Fig. 3S). Taken together, our data suggest that Lar and Sns interact in *cis* in motor neurons to regulate NMJ development.

### Lar and Sns reporters do not colocalize in the larval mushroom body

Next, we analyzed Lar and Sns expression in the larval brain, specifically focusing on the MB, as Lar has been shown to be required for proper development of the larval MB (Kurusu and Zinn, 2008). Lar was shown to be expressed in Kenyon cells (KCs), the principal cells of the MB, using antibody staining against LAR protein. Here, we confirm that Lar is expressed in larval KCs using Lar>GFP (Fig. 4A). Lar>GFP is also seen in MB axons that compose the two lobes of the larval MB, the dorsal lobe and the medial lobe (Figs. 4B, B’). A confocal *z*-projection through the entire larval MB is shown in Figure 4B, B’. The MB lobes are visualized using an antibody against Fasciclin II (FasII), which specifically labels the MB neuropil (Figure 4B). A single optical slice shows that Lar>GFP labels both the dorsal (arrow) and the medial lobes of the larval MB (Figure 4C’).

**Figure 4:**
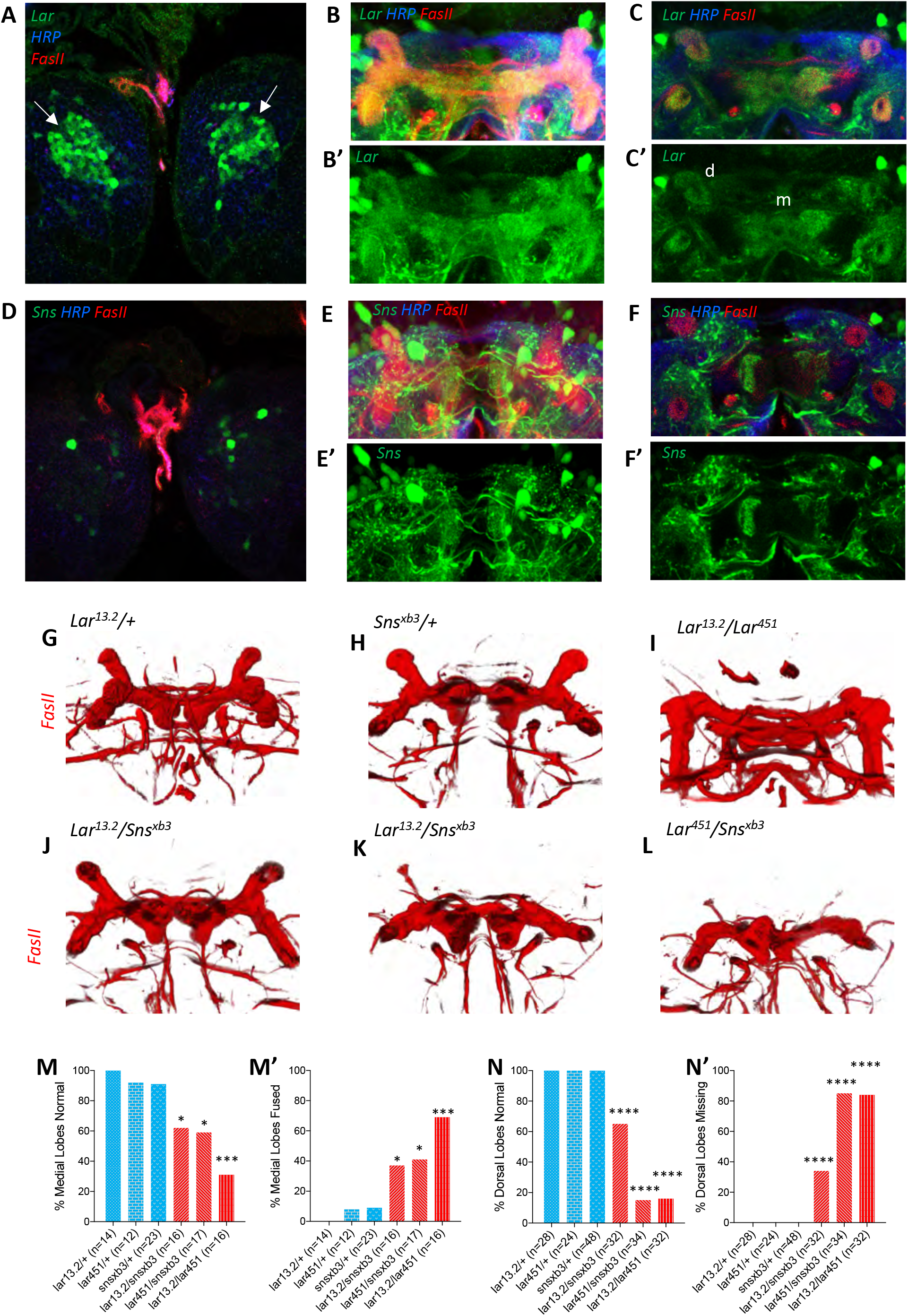
Lar and Sns act in different neurons to control MB dorsal and medial lobe development. (A-F’) Lar>GFP and Sns>GFP expression in the larval brain. Brains were triple-stained for Lar>GFP (green), FasII (red), and anti-HRP (blue). Anti-FasII labels the MB neuropil; anti-HRP labels neuronal membranes. (A) Lar>GFP expression in Kenyon cells (green). (B-B’) Projection of confocal slices through the entire larval MB showing Lar expression in the MB neuropil. (C, C’) Single optical slice showing Lar expression in the medial (m) and dorsal (d) lobes of the MB. (D) There is no Sns>GFP expression seen in Kenyon cells. (E, E’) Projection of confocal slices through the entire mushroom body showing no overlap between Sns>GFP and the MB neuropil labeled by FasII. (F, F’) Single optical slice through the mushroom body showing no Sns>GFP expression in the MB neuropil. (G-L) Third instar larval MBs visualized with FasII staining. 3D reconstructions of confocal stacks using Imaris software are shown. (G) and (H) have normal MBs. (J) has a medial lobe fusion phenotype. (K) has missing dorsal lobes. (L) has missing dorsal lobes and medial lobe fusion. (M-N’) Quantification of MB phenotypes in heterozygote controls (blue), *Lar* mutants (red), and *Lar/sns* transhets (red). In (M) and (N), the % of normal MBs are shown; in (M’) and (N’), the % of MBs with the phenotype are shown. (M, M’) Medial lobe fusion phenotype, (N, N’) dorsal lobe branching defect. Data were analyzed using Fisher’s Exact Test and each genotype was compared to every other genotype. **** p<0.0001; *** p<0.001; * p<0.05.

Sns>GFP was not detected in KCs (Fig. 4D), but was seen in many other neurons in the larval central brain. No Sns reporter expression could be seen in either the dorsal or the medial lobes of the MB (Figs. 4E, E’). A single optical slice of the MB lobes shows no Sns>GFP expression in the MB neuropil (Figs. 4F, F’). In order to determine if Sns is expressed in neurons postsynaptic to MB axons, we used a dendrite-specific marker (UAS-Drep2) (Andlauer et al., 2014) to label dendrites of Sns-expressing neurons. We observed dendrites of Sns-expressing neurons enveloping the dorsal lobes of the mushroom body (Supp. Fig. 1), indicating that Sns-expressing neurons are postsynaptic to Lar-expressing KCs. Next, we asked whether Sns>GFP was expressed in glial cells in addition to neurons. We performed immunostaining using Sns>GFP and anti-Repo to label glial cells. We did not see any co-localization between GFP and Repo (Supp. Fig. 1), showing that Sns is expressed only in neurons.

### Lar and Sns genetically interact to shape the formation of the larval mushroom body

Our group has previously shown that Lar is required for the formation of the larval MB, with Lar mutants displaying two distinct defects in the third instar MB. In WT animals, KC axons bifurcate and form two separate neuropil lobes, the medial lobe and the dorsal lobe. Axons in the medial lobe stop at the midline and do not cross over to the contralateral side. In *Lar* mutants, these medial lobe axons fail to stop at the midline, instead crossing over to the contralateral side and forming a fused medial lobe across the midline. A second defect is seen in *Lar* mutants in which KC axons do not branch properly, resulting in reduced or absent dorsal lobes (Kurusu and Zinn, 2008). Ligand(s) involved in Lar’s actions in the larval MB have not been identified. We investigated whether Sns is required for Lar’s role in the development of the larval MB by phenotypic analysis of *Lar/sns* transhets.

We analyzed the medial and dorsal lobes in *Lar/sns* transhets and *Lar* mutants, along with their respective heterozygote controls. FasII antibody staining specifically labels both medial and dorsal lobes. We analyzed 3D reconstructions of FasII-stained larval MBs to visualize the lobes in their entirety. Each optical section of confocal *z*-stacks through the MBs was analyzed for the medial lobe fusion phenotype. Supp. Fig. 3 shows single optical slices with medial lobe axons either intact or crossing the midline.

For clarity, data on medial lobe and dorsal lobe phenotypes is displayed in two types of bar graphs, showing % of MBs (or animals) without the phenotype (*e.g.,* Fig. 4M), and % of MBs with the phenotype (*e.g.,* Fig. 4M’). Heterozygous control animals did not show any abnormal phenotypes in the larval MB, with largely normal medial and dorsal lobes (Figs. 4G-H, Supp. Fig. 3). However, *Lar*^*13.2*^/*sns*^*xb3*^ transhets displayed fused medial lobes in 37% of animals, and *Lar*^*451*^/*sns*^*xb3*^ animals had fused medial lobes in 41% of animals. *Lar* mutants displayed fused medial lobes in 69% of animals (Figs. 4I-L, M, M’). We also observed dorsal lobe phenotypes in *Lar/sns* transhets, with 34% of dorsal lobes either reduced or missing in *Lar*^*13.2*^/*sns*^*xb3*^ transhets and 85% of dorsal lobes reduced or missing in *Lar^451^/sns^xb3^* animals. 84% of *Lar* mutants displayed this branching defect, lacking dorsal lobes (Figs. 4I, K, L, N, N’). We did not observe a strong correlation between the two phenotypes occurring in the same animal. An animal with a fused medial lobe did not necessarily also display reduced or absent dorsal lobes.

We next performed pan-neuronal RNAi knockdown for Lar and Sns using the two RNAi lines for each gene to investigate whether knocking down each gene individually also results in the MB abnormalities seen in *Lar/sns* transhets and *Lar* mutants. Knocking down Sns resulted in medial lobe fusion in 58% of animals using the KK line and 42% of animals using the GD line (Supp. Fig. 3). *Lar* RNAi knock-down resulted in 100% of animals showing fused medial lobes with the stronger RNAi line (HMS02186) and 54% of animals with fused medial lobes using the second RNAi line (HMS00822). We observed weaker penetrances of the branching defect with RNAi, with only 11% reduced or missing dorsal lobes upon Sns knockdown with the KK line and 21% reduced or missing dorsal lobes with the GD line (Supp. Fig. 3). Similarly, only 27% of dorsal lobes were reduced or missing upon Lar knock-down with the second RNAi line (HMS00822), while the stronger Lar RNAi line displayed 61% penetrance (Supp. Fig. 3). Thus, knocking down Lar and Sns causes similar developmental abnormalities as those seen in transhets and *Lar* mutants.

These data show that Lar interacts with Sns to regulate the formation of the larval MB. The Lar-Sns interaction in this context is likely to be in *trans*, as we do not observe any Sns>GFP expression in larval KCs or in the MB lobes. Lar>GFP, on the other hand, is strongly expressed in larval KCs and the reporter is visible in both the medial and the dorsal lobes. We observe dendrites of Sns-expressing neurons encircling the dorsal lobes of the mushroom body, indicating that Sns might be in neurons that are postsynaptic to Lar-expressing KC axons.

### Expression patterns of Lar and Sns in the pupal and adult mushroom body

To further clarify the relationships between the Lar and Sns expression patterns, we examined Lar>GFP and Sns>GFP in the adult MB. Adult KCs can be broadly classified into three neuronal types based on the lobes they innervate. γ neurons are born before the third instar larval stage and form the adult γ lobe. α’/β’ neurons are born during the late third instar larval stage and form the α’ and the β’ lobes, which project dorsally and medially respectively. α/β neurons are born during early pupal stages and form the α/β lobes, which project dorsally and medially, parallel to the α’ and β’ lobes. α/β lobes can be visualized using FasII antibody staining, while the γ and α’/β’ lobes can be visualized using Trio antibody staining.

We performed immunostaining for either FasII or Trio combined with anti-GFP staining to label either Lar>GFP and Sns>GFP adult brains. We found strong Lar expression in the α/β lobes of the adult MB, with strong co-localization of FasII with Lar>GFP (Figs. 5A, A’). Single optical slices through the adult MB show strong Lar>GFP expression in both the α and β lobes (Figs. 5B, B’). However, we did not observe any expression in the α’/β’ lobes, which were labeled by Trio antibody staining (Figs. 5C, C’). Single optical slices showed clear Lar>GFP expression in α/β lobes, but no detectable expression in α’/β’ lobes (Figs. 5D, D’). We observed weak Sns>GFP expression in the α/β lobes. Figs. 5E, E’ show a single optical slice. There was no co-localization of Sns>GFP and Trio antibody staining (Figs. 5F, F’). Thus, neither Lar or Sns is detectably expressed in the adult α’/β’ lobes.

**Figure 5:**
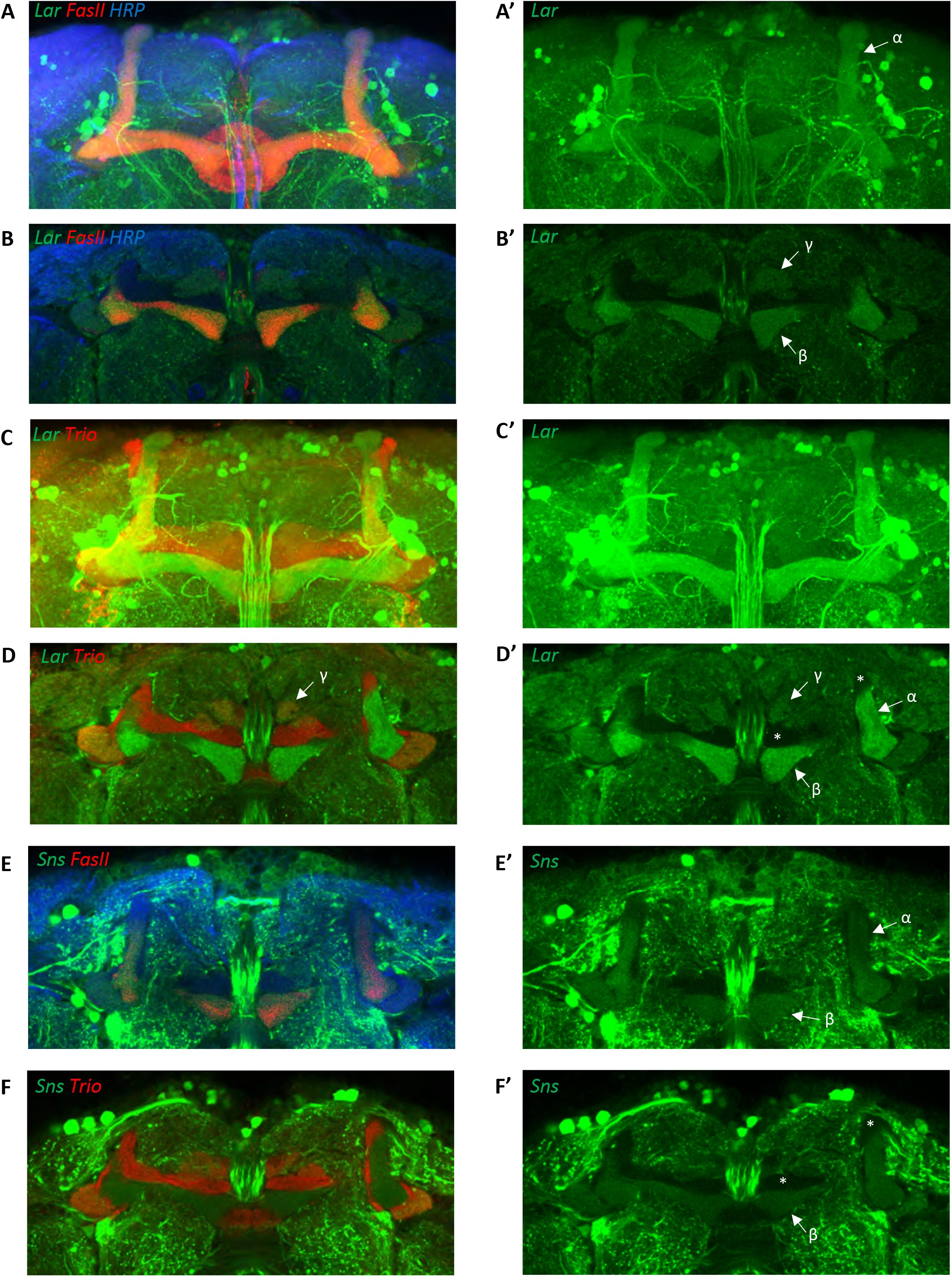
Lar and Sns expression in the adult MB. Confocal projections and single optical slices showing Lar>GFP and Sns>GFP (A, B) Projection of confocal slices showing the entire adult mushroom body, triple-stained for GFP (green), FasII (red) and anti-HRP (blue). (A’, B’) GFP signals only. (A) FasII labels the α and β lobes of the adult MB and is colocalized with Lar>GFP. (B) Single optical slice from the adult MB showing strong Lar>GFP expression in the α and β lobes, co-localizing with FasII staining. (C, C’) Projection of confocal slices showing the entire adult MB, double-stained for GFP (green) and Trio (red). Trio labels the γ, α’ and β’ lobes of the adult MB. (D, D’) Single optical slice showing no detectable Lar>GFP expression in the Trio-expressing α’ and β’ lobes. Weak expression is seen in the γ lobe (B’). (E, E’) Single optical slice showing faint Sns expression in the α and β lobes (Sns, green; FasII, red). (F, F’) Single optical slice showing no Sns expression in the γ, α’ and β’ lobes (Sns, green; *Trio*, red).

We next investigated the developmental profile of Lar and Sns expression in the MB during pupal stages. We analyzed Lar>GFP and Sns>GFP expression in the MB in 24APF, 40APF and 72APF pupal brains. We observed very little Lar>GFP in the 24APF MB (Supp. Figs. 4A, A’). We did see Lar expression on growth cones of α/β KC axons near the midline in the β lobes (Supp. Figs. 4B, B’, arrows). At 40APF, moderate Lar>GFP expression can be seen in both α/β lobes as well as α’/β’ lobes. A single optical slice shows clear Lar>GFP expression in α/β and α’/β’ lobes (Supp. Figs. 4F, F’) Lar expression peaks at 72APF in the MB, with strong Lar>GFP expression seen in all lobes, the γ lobe, α/β lobes and α’/β’ lobes. Single optical slices show that Lar>GFP is expressed in both α/β lobes and α’/β’ lobes at 72APF (Supp. Figs. 4J-K’). Thus, Lar expression is high during the time period of active MB axonal outgrowth and synaptic targeting.

Sns>GFP is expressed at high levels in several neuronal populations in the central brain at all three pupal stages. However, we did not observe any detectable Sns>GFP expression in either the α/β lobes or the α’/β’ lobes in the 24APF and 40APF MB (Supp. Fig. 4). Weak Sns>GFP expression can be seen in α/β lobes, but not α’/β’ lobes, at 72APF. Kenyon cell axons are born sequentially, with newer-born axons in the center of the lobes pushing the older axons to the outer edges of the lobes. Sns>GFP seemed to be selectively expressed by older axons in the α and the β lobes as we observed hollow cores with no GFP labeling in both lobes (Supp. Fig. 4M-N’, arrows and asterisks in hollow cores).

In addition to examining the MB, we also characterized Lar and Sns expression in the pupal and adult antennal lobes, which contain projection neurons (PNs) that synapse onto KCs. As in the MB, we observed the highest expression at 72 hr. APF, when Sns>GFP and Lar>GFP labeled specific glomeruli (Supp. Fig. 5). This likely represents PN expression, since it has been demonstrated that Lar and Sns are enriched in PNs (Li et al., 2020).

### Lar and Sns genetically interact to regulate morphogenesis of α/β and α’/β’ lobes of the adult mushroom body

We have shown that Lar and Sns genetically interact to regulate the development of larval MB lobes. Since Lar is expressed at high levels in the developing pupal MB, we investigated whether Lar and Sns play a role in the morphogenesis of the adult MB lobes as well. We used FasII immunostaining to visualize the α and β lobes and Trio immunostaining to visualize the γ, *α*’, and *β*’ lobes.

Heterozygote controls (*Lar^13.2^*/+, *Lar*^*451*^/+, and *sns^xb3^*/+) animals all have normal α and β lobes (Figs. 6A-C, G-H’). Note that the β lobes in these controls end well before the midline (asterisks in Figs. 6A-C). *Lar*^*13.2*^/*sns*^*xb3*^, *Lar*^*451*^/*sns*^*xb3*^ and *Lar*^*13.2*^/*Lar*^*451*^ animals showed 75%, 79% and 93% missing α lobes, respectively. *Lar/sns* transhets and *Lar* mutants also displayed midline crossing of β lobe axons, with most β lobe axons crossing the midline, creating a fused β lobe, instead of two separate lobes (Figs. 6D-H’). We noted that the thickness of these fused β lobes was significantly greater than the normal unfused β lobes seen in control animals.

**Figure 6:**
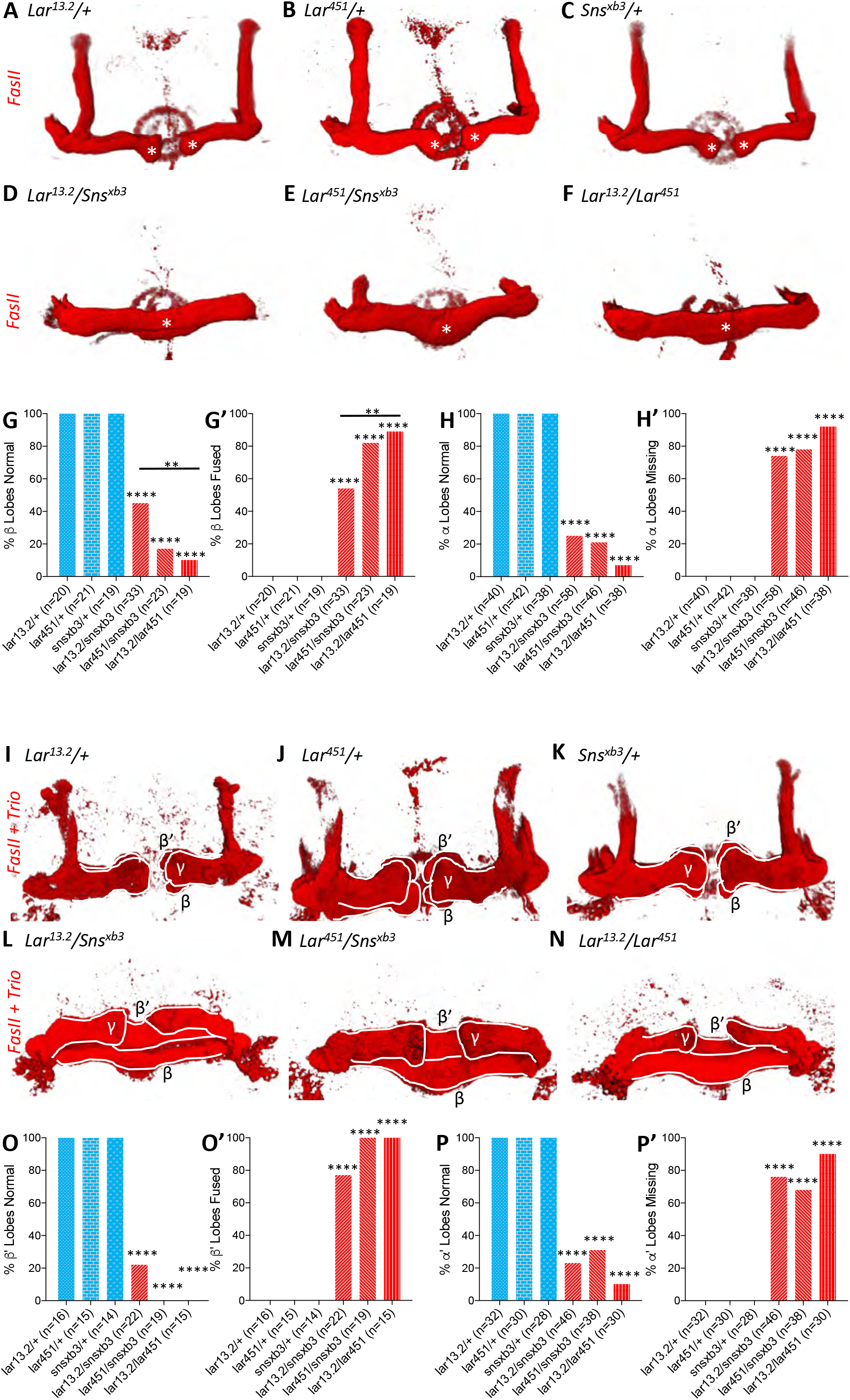
Lar and Sns are required for normal lobe development in the adult MB. (A-F) 3D reconstructions of confocal stacks from anti-FasII stained adult brains using Imaris software. (A-C) Heterozygote controls showing normal α and β lobes of the adult MB. Asterisks show the ends of normal β lobes, which stop short of the midline and remain separated. (D-F) *Lar/sns* transheterozygotes and *Lar* mutants, showing abnormal MB architecture, with missing α lobes and β lobes fused across the midline. (G, G’) Quantification of β γlobe midline fusion phenotype. Heterozygote controls (blue) show completely normal unfused β lobes. *Lar/sns* transheterozygotes and *Lar* mutants (red) have fused β lobes. (H, H’) Quantification of α γlobe branching defect. Heterozygote controls (blue) have intact α lobes, while *Lar/sns* transhets and *Lar* mutants (red) have missing α lobes. In (G) and (H), the % of normal MBs are shown; in (G’) and (H’), the % of MBs with the phenotype are shown. (I-N) 3D reconstructions of confocal stacks from adult brains stained with *FasII* and *Trio* antibody to visualize the entire MB with all lobes. (I-K) Heterozygote controls show normal mushroom body lobes. (L-N) *Lar/sns* transheterozygotes and *Lar* mutants show abnormal mushroom body architecture, with fused β and β’ lobes and missing α and α’ lobes. (O, O’) Quantification of β’ lobe midline fusion phenotype showing normal β’ lobes in heterozygote controls (blue) and almost completely fused β’ lobes in *Lar/sns* transheterozygotes and *Lar* mutants (red). (P, P’) Quantification of α’ lobe branching defect. Heterozygote controls (blue) have completely normal α’ lobes while *Lar/sns* transhets and *Lar* mutants (red) are missing most α’ lobes. In (O) and (P), the % of normal MBs are shown; in (O’) and (P’), the % of MBs with the phenotype are shown. Data were analyzed using Fisher’s Exact Test and each genotype was compared to every other genotype. **** p<0.0001; ** p<0.01.

We then performed combined FasII and Trio immunostaining to visualize all the lobes of the MB. γ lobes in all animals were normal and showed no phenotype in either *Lar/sns* transhets or *Lar* nulls (Figs. 6I-N). Similar to α lobes, we observed normal α’ lobes in *Lar* and *Sns* heterozygote control animals (Figs. 6I-K). *Lar/sns* transhets and *Lar* nulls displayed highly abnormal α’ lobes, with most animals missing one or more α’ lobes (Fig. 6L-N, P, P’). Heterozygote control animals showed completely normal β’ lobes (Figs. 6I-K, O, O’). However, β’ lobes displayed midline crossing resulting in a single fused β’ lobe in most or all *Lar/sns* transhets and *Lar* null animals (Fig. 6L-N, P, P’). Thus, the *Lar*, *sns*, and *Lar/sns* phenotypes were even stronger in β’ lobes than in β lobes. Since we do not detect Sns expression in pupal or adult α’/β’ KCs, Sns must be acting in some other neuronal type to regulate α’/β’ lobe development, and the relevant Lar-Sns interaction is likely to be in *trans*, mediated by interactions between Lar and Sns expressed on apposed neurons.

### Lar and Sns expression in the pupal and adult optic lobe

*Lar* mutants have well-characterized phenotypes in the OL, where Lar is required for innervation of medulla layer M6 by R7 photoreceptor axons (Clandinin et al., 2001; Hakeda-Suzuki et al., 2017; Hofmeyer and Treisman, 2009; Maurel-Zaffran et al., 2001). To analyze whether Sns is involved in this Lar function as well, we characterized expression of Lar>GFP and Sns>GFP reporters in the OL. Interestingly, although Lar has been studied in the OL for years, there has been no characterization of Lar expression based on a GAL4 reporter. Lar antibody staining is not informative about cell-specific expression patterns, because the antibody uniformly labels the neuropil and does not stain cell bodies (Maurel-Zaffran et al., 2001).

Both Lar>GFP and Sns>GFP were strongly expressed in the adult OL, with expression seen in all neuropils. We analyzed expression in one and 7 day old adult flies, and while the expression pattern of Lar remained the same, we found that Lar>GFP expression levels were reduced in 7 day old flies compared to one day old flies (Figs. 7A-C’). Strong Lar>GFP expression was seen in L1 lamina neurons. The cell bodies of L1 neurons are in the lamina, and they send their axonal/dendritic projections to the M1 and M5 layers of the medulla. We observed strong Lar>GFP expression in L1 cell bodies and in M1 and M5 layers of the medulla in one day old flies (Figs. 7B, B’), but this had weakened in L1 cell bodies in 7 day old flies (Figs. 7A, A’). We still observed punctae of GFP staining in M1 and M5 layers of the medulla, but the expression level was much lower than in one day old flies (Fig. 7A’’). We did not observe detectable Lar>GFP expression in any photoreceptors in the adult optic lobe. Lar>GFP expression was also seen in two layers of the lobula neuropil (Figs. 7A, A’).

**Figure 7:**
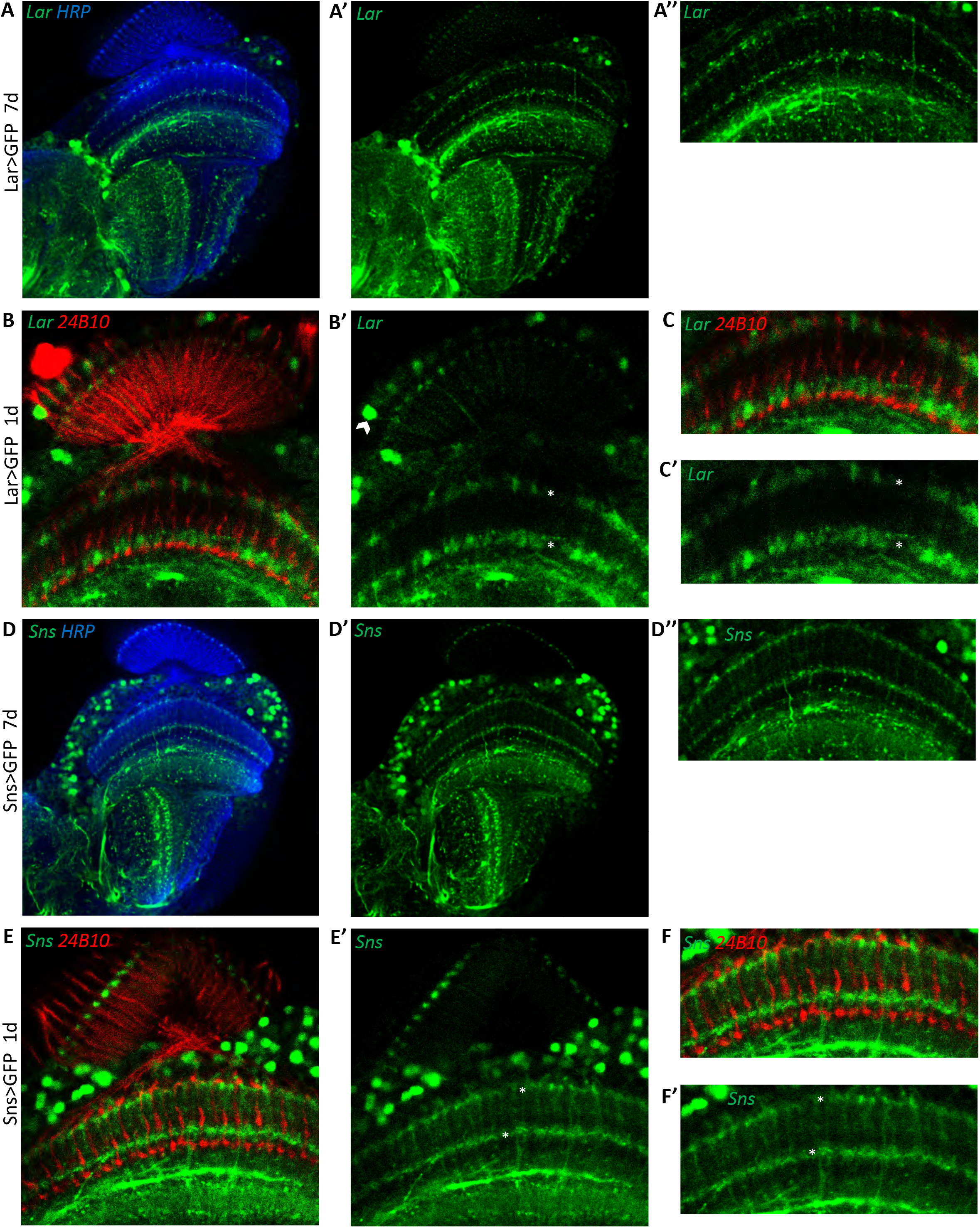
Expression of Lar and Sns in the adult OL. Confocal projections and single optical slices showing Lar>GFP and Sns>GFP expression in the OL. Brains were double-stained for GFP (green) and either anti-HRP (blue) or Chaoptin (24B10; red), which labels photoreceptor axons. (A-A’’) Single optical slice showing Lar expression (green) in seven-day old adult fly optic lobe. (A’’) is a higher magnification view of (A’). (B-C’) Single optical slice showing Lar>GFP expression (green) in one-day old fly optic lobe). Stronger Lar>GFP expression is seen in one-day old optic lobe in layers M1 and M5 (asterisks in B’ and C’) of the medulla. (C) and (C’) are higher-magnification views of (B) and (B’), respectively. Lar>GFP expression becomes more punctate in seven-day old optic lobe. (D-D’’) Single optical slice showing Sns>GFP expression (green) in seven-day old fly optic lobe. (E-F’) Single optical slice showing Sns>GFP expression (green) in a one-day old fly optic lobe). Sns>GFP expression stays the same between one and seven-day old optic lobe. Strong Sns>GFP expression is seen in layers M1 and M5 of the medulla. Sns>GFP does not appear to colocalize with 24B10 in (F) and (F’), suggesting that it is not in R7 or R8 photoreceptors (see Supplemental Fig. 7).

Sns>GFP expression was seen in numerous cells in the medullary cortex (Figs. 7D-E’). Unlike Lar, Sns>GFP expression levels did not decrease significantly in 7 day old flies (compare Figs. D’ and E’). We observed Sns>GFP expression in spots at the top of the lamina neuropil, and also in the M1 and M5 layers of the medulla. This matches the arborization pattern of C2 neurons, which are synaptic partners of L1 neurons (Supp. Fig. 7C-C’’). C2 neurons are the only neurons with this dot-like pattern of endings, one at the top of each lamina cartridge (see Tuthill et al., 2013). We used a dendrite-specific marker (Drep2) to label dendrites of Sns-T2A-GAL4-expressing neurons in the adult OL and found that the Sns expression seen in lamina and the M1 and M5 layers of the medulla at least partly represents postsynaptic elements (Supp. Fig. 7A, A’). C2 neurons are bidirectionally connected to L1 neurons in M1 and M5 (Takemura et al., 2013; Takemura et al., 2015). Thus, Lar and Sns might interact in trans to regulate development of the L1-C2 circuit. Finally, we also observed strong Sns>GFP expression in the proximal layers of the medulla, as well as in two layers of the lobula (Figs. 7D, D’).

We also analyzed Lar and Sns expression in the developing pupal OL at 24, 40 and 72 hr. APF. We observed strong Lar>GFP expression at both 40 and 72 hr. APF in the L1 lamina neuron cell bodies (Supp. Fig. 6C, C’, G, G’). Lar>GFP was also expressed at high levels in the M1 and M5 layers of the medulla where L1 neurons arborize, with higher expression seen in 72 hr. APF pupal brains. This is consistent with results of sequencing of pupal lamina neuron mRNA, which showed that *Lar* is expressed at very high levels in pupal L1 (Tan et al., 2015). We also found that Lar>GFP was expressed in several photoreceptor cell bodies in the 24APF retina (Supp. Figs. 7B, B’). Stronger expression was seen in R7 and/or R8 photoreceptors, which lie in the center of each ommatidium. Lar>GFP was also expressed in at least one other outer photoreceptor.

Sns>GFP was expressed at high levels in the developing 40 and 72 hr. APF OL (Supp. Fig. 6). Many neurons expressed Sns>GFP in the 40 and 72 hr. APF medullary cortex. Strong Sns expression was seen in both the proximal and distal layers of the medulla at 40 hr. APF. By 72 hr. APF, Sns expression could be seen in dots at the top of the lamina, which are likely to be C2 endings (Supp. Fig. 7C, C’), as well as in the M1 and M5 layers of the medulla (Supp. Fig. 6J-J’). We did not detect Sns>GFP expression in any photoreceptors at any pupal stage.

### Lar and Sns interact to regulate R7 photoreceptor axon targeting

Lar has been shown to be required for R7 axon targeting to the M6 layer of the medulla. In *Lar* mutants, R7 axons initially project to the correct M6 layer, but later retract to the M3 layer during mid-pupal stages (Clandinin et al., 2001; Maurel-Zaffran et al., 2001). We analyzed R7 photoreceptor axon targeting in *Lar/Sns* transhets and *Lar* mutants using Chaoptin (Chp) immunostaining, which labels all photoreceptors. In the adult medulla, R7 axon endings can be clearly seen in the M6 layer by Chp (24B10) staining. Heterozygote controls of *Lar*^*13.2*^, *Lar*^*451*^ and *sns*^*xb3*^ have R7 axons that terminate in the appropriate M6 layer (Figs. 8A-C, white outlines) and have normal R7 terminal morphology. In both *Lar/sns* transhets (*Lar*^*13.2*^/*sns*^*xb3*^ and *Lar*^*451*^/*sns*^*xb3*^), R7 axons failed to terminate in the appropriate M6 layer, instead retracting to the M3 layer (Figs. 8D, E). In *Lar*^*13.2*^/*sns*^*xb3*^ transhets, 71% of R7 axons were retracted, and in *Lar*^*451*^/*sns*^*xb3*^ transhets, 63% of R7 axons were retracted (Fig. 8G). *Lar^13.2^/Lar^451^* mutants had similar phenotypes, with 88% of R7 axons retracted to the M3 layer (Figs. 8F, G). In addition, we also observed that the R7 axons that did innervate the M6 layer had abnormal terminal morphologies. Normal R7 terminals have a rounded bouton-like appearance (Fig. 8H, arrow). In *Lar/sns* transhets and *Lar* mutants, R7 axon terminals have a spear-like appearance with thin axon terminals (Fig. 8I, J, arrows). Previous studies have shown that Lar acts cell-autonomously in R7 neurons to regulate R7 axon targeting, and we found that Lar>GFP is expressed in R7 and/or R8 photoreceptors during pupal development (Supp. Fig. 8B, B). However, there was no Sns>GFP expression in photoreceptors. Therefore, Sns must be acting in some other neuronal type to regulate R7 axon targeting. The M5 labeling by Sns>GFP reflects Sns expression in C2, and perhaps in other medulla neurons that arborize in the layer immediately above that targeted by R7 axons. Perhaps interactions between Lar in R7s and Sns in this neuron(s) facilitate R7 axon adhesion to the M6 layer and prevent retraction.

**Figure 8:**
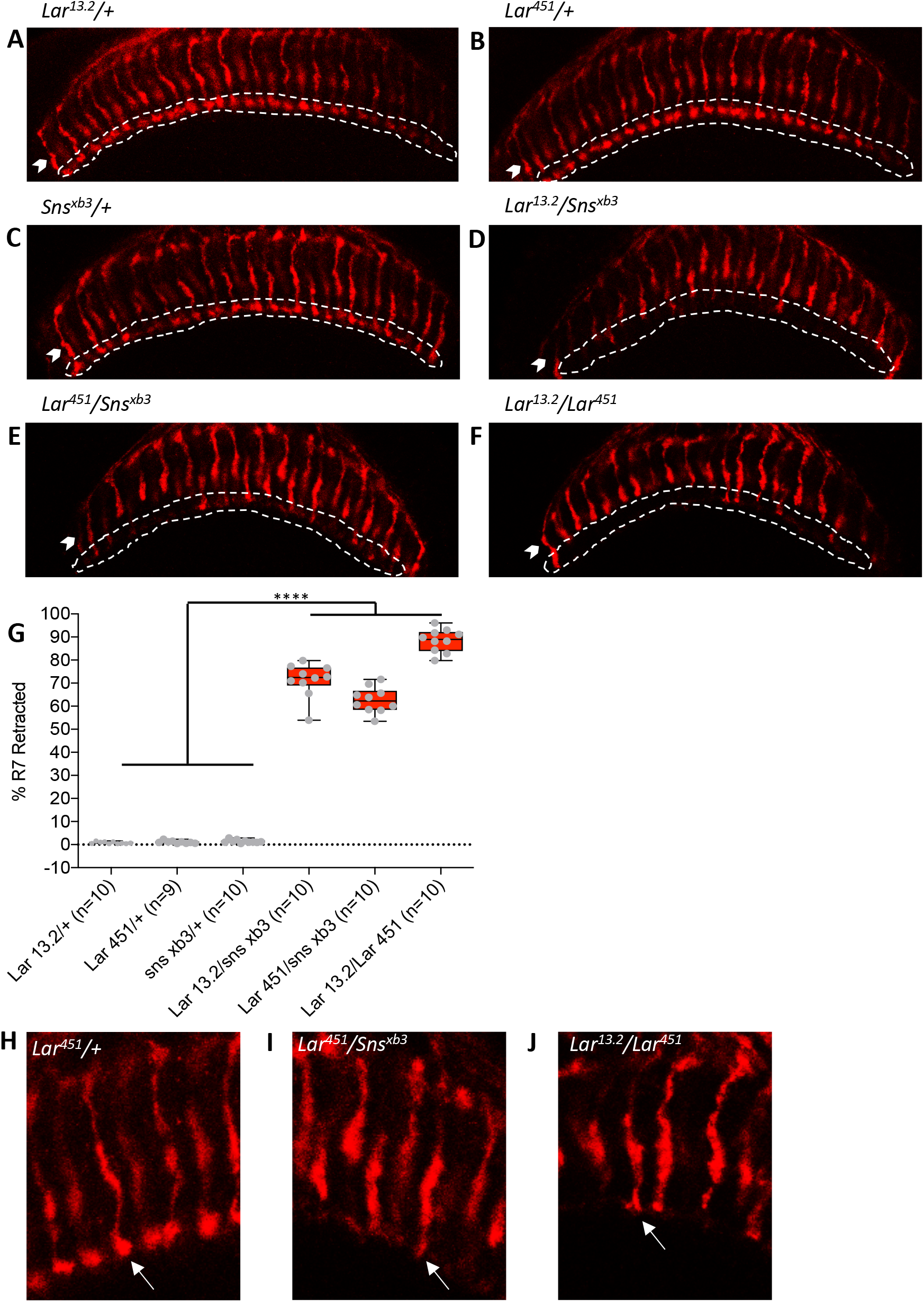
R7 photoreceptors have identical targeting defects in *Lar* mutants and *Lar/sns* transheterozygotes. (A-F) Single optical slices of adult OLs showing R7 and R8 photoreceptors labeled for Chaoptin (24B10, red). R7 photoreceptor axons end in the deeper M6 layer of the medulla (outlined in white), while R8 axons end in M3 layer (arrowheads). (A-C) Heterozygote controls showing normal R7 targeting in the M6 layer. (D-F) *Lar/sns* transheterozygotes and *Lar* mutants, showing abnormal R7 targeting, with most R7 axons retracting to the M3 layer. (G) Quantification of R7 axon retractions in control and mutant animals. R7 axons were counted in at least 10 optical slices per optic lobe. Each data point is the average of 10-12 optical slices per optic lobe. The data were analyzed using One-Way ANOVA followed by Tukey’s post-hoc correction. **** p<0.0001. (H-J) Single optical slices showing the morphology of R7 terminals in *Lar^451^*/+ control, *Lar^451^/sns^xb3^* transheterozygotes, and *Lar* mutants. Control animals show normal rounded bouton-shaped R7 terminals (H, arrow). Some R7 terminals that do not retract and stay in the M6 layer have abnormal R7 terminal morphologies, with thin and spear-shaped terminals (I, J, arrows).

## Discussion

In this paper, we provide evidence that Sns, a large cell-surface protein known for its roles in muscle fusion and cell patterning, is a ligand for the Lar RPTP that is required for Lar function during neural development. We identified Sns as a Lar ligand by employing an ectopic ligand binding screen in embryos. This screen had previously identified Sas as a ligand for PTP10D (Lee et al., 2013). Lar binds to Sns both in forward (Lar-AP_5_ staining of embryos overexpressing Sns) and reverse (Sns-AP_5_ staining of Lar-expressing embryos) binding experiments (Fig. 1). Lar and Sns also bind to each other *in vitro* in a modified ELISA assay. This interaction appears to be evolutionarily conserved, because at least two of the three human Lar orthologs, PTPRF and PTPRD, bind to the Sns ortholog Nephrin, as well as to Sns, in embryos and *in vitro* (Fig. 1).

Having shown that Lar and Sns are binding partners, we then examined whether Sns is required for Lar function *in vivo* by assessing the phenotypes of transheterozygous animals lacking one copy of each gene. We also examined Lar and Sns expression to determine whether Lar and Sns are likely to interact in *cis* (on the same neuron) or in *trans* (between neurons). Lar is known to regulate NMJ morphogenesis, and we showed that Sns is required for this function and that the two proteins are exclusively expressed in motor neurons and therefore work together in *cis* (Figs. 2–3).

Lar is also required for larval MB development and is expressed in MB KCs, while Sns is not expressed in the MB. Sns is required for Lar’s functions in midline stopping of KC axons and in KC axon branching to form the dorsal lobe (Fig. 4, Supp. Fig. 3). Lar and Sns also act in *trans* during pupal/adult MB development, and transheterozygous phenotypes there are even stronger (Figs. 5, 6, Supp. Fig. 4). Finally, Lar and Sns are expressed in different sets of neurons in the pupal OL (Fig. 7, Supp. Fig. 6). *Lar* acts cell-autonomously in R7 neurons to facilitate innervation of the M6 medulla layer by R7 axons (Maurel-Zaffran et al., 2001). However, Sns is not expressed in photoreceptors. Sns on medulla neurons may interact in *trans* with Lar on R7 growth cones to facilitate innervation of the M6 layer (Fig. 8).

### Lar and Sns act in *cis* to regulate NMJ morphogenesis

Lar and Sns are expressed in the same motor neurons. Lar as well as Sns expression was seen in several 1b type of motor neurons, including the 1b motor neuron that innervates muscles 7 and 6 (7/6). Both Lar and Sns were also expressed in the two 1s motor neurons, MNISN-1s and MNISNb/d-1s. Each 1s motor neuron arborizes onto several muscles in a field. We found that the interaction between Lar and Sns is required for NMJ development at the 7/6 NMJ. Both 1b and 1s types of arbors at the 7/6 NMJ were abnormal in *Lar/sns* transhets. This phenotype was indistinguishable from that of *Lar* mutants. We could not analyze this phenotype in *sns* mutants, since *sns* null mutants do not survive beyond early first instar larval stage. Instead, we performed RNAi experiments using a neuronal-specific driver and knocked down both Lar and Sns. Again, we saw that knocking down Lar and Sns resulted in the same NMJ phenotypes. The lack of Lar and Sns expression in muscles, combined with their expression in motor neurons that innervate the 7/6 NMJ, suggest that Lar and Sns act in *cis* in motor neurons to regulate NMJ development (Figs. 2, 3, Supp. Fig. 2).

A possible mechanism of Lar and Sns’s effects on NMJ development could be by regulating actin polymerization. SYG-2 and SYG-1, the *C. elegans* orthologs of Sns and Kirre, respectively, have been shown to regulate presynaptic assembly and branch formation on HSN synapses by directly affecting F-actin assembly (Chia et al., 2014). During myoblast fusion, Sns interacts with several proteins including Wiskott-Aldrich syndrome protein (WASP), Solitary (Sltr)/dWIP, and the GTPase Rac. Sns signaling ultimately results in reorganization of the actin cytoskeleton resulting in myoblast fusion between founder cells and fusion competent myoblasts (Sens et al., 2010). At *Drosophila* NMJs, bouton assembly and terminal branching are dependent upon F-actin assembly (Koch et al., 2014). Several studies have demonstrated a link between terminal arborization and synapse assembly (Chia et al., 2014; Koch et al., 2014). We have shown that the Lar-Sns interaction is required for both NMJ terminal arborization as well as synapse assembly. Since Sns has been shown to regulate F-actin dynamics during its actions in myoblast fusion, we hypothesize that the Lar-Sns interaction may also regulates F-actin dynamics at the developing NMJ. Perhaps Lar and Sns form a complex on the neuronal cell surface, and dephosphorylation by Lar directly or indirectly regulates some of the proteins of the actin cytoskeleton that interact with Sns. Two key substrates of Lar, the Abl tyrosine kinase and Ena, are regulators of actin assembly (Johnson and Van Vactor, 2003). Moreover, the cytoplasmic domain of Sns is phosphorylated at multiple tyrosines (Kocherlakota et al., 2008). Sns could thus be a direct target for dephosphorylation by Lar.

### *Trans* interactions between Lar and Sns regulate mushroom body axon guidance

Our group previously showed that Lar is required in the developing third instar larval MB for KC axon guidance (Kurusu and Zinn, 2008). *Lar* mutants display abnormal axon guidance in KC axons, resulting in two distinct phenotypes: first, the KC axons fail to stop at the midline and instead extend into the contralateral medial lobe, resulting in a fused medial lobe. Second, KC axons fail to branch at the branching point near the peduncle, resulting in a reduced or completely absent dorsal lobe (Kurusu and Zinn, 2008). The Lar>GFP reporter is expressed in larval, pupal, and adult KCs (Fig. 4, Supp. Fig. 3).

Here we show that Sns acts together with Lar to regulate axonal midline stopping and branching in the larval as well as in the pupal/adult MB (Figs. 4–6, Supp. Fig. 3). *Lar/sns* transhets and Sns knockdown animals have the same larval phenotypes as *Lar* mutants. The Sns reporter is not expressed in larval KCs (Fig. 4, Supp. Figs. 1, 3). In the adult MB, both the α/β and α’/β’ lobes are strongly affected in *Lar/sns* transhets and *Lar* mutants. The Sns reporter is not expressed in α’/β’ KCs at any time, and exhibits weak α/β KC expression only after lobe development has taken place (Figs. 5, 6, Supp. Fig. 4). These data show that Sns is likely to interact in *trans* with Lar in the MB system. We observed that Sns is expressed in larval neurons (MBONs) that have postsynaptic elements apposed to KC axons (Supp. Fig. 1). Perhaps Sns expressed on these neurons acts as a guidance cue for the KC axons.

### Sns is also required in *trans* for Lar’s roles in R7 photoreceptor axon targeting

Another known role of Lar is in R7 photoreceptor axon targeting. *Lar* mutants display aberrant R7 targeting in the adult optic lobe, with most R7 axon terminals retracted to the M3 layer of the medulla instead of the correct M6 layer (Clandinin et al., 2001; Hakeda-Suzuki et al., 2017; Hofmeyer and Treisman, 2009; Maurel-Zaffran et al., 2001). The ligand(s) responsible for Lar’s actions in R7 axon targeting have not been identified. Previous research showed that the last three FNIII domains of Lar were required for R7 targeting, and that the Ig domains of Lar were not involved (Hofmeyer and Treisman, 2009). Syndecan binds to the Ig domains of Lar (Fox and Zinn, 2005; Johnson et al., 2006), so it could be ruled out as the Lar ligand responsible for mediating R7 targeting.

More than 70% of R7 terminals in *Lar/sns* transhets show retraction to the M3 layer. Previous research has shown that Lar acts cell autonomously in R7 photoreceptors to regulate targeting (Maurel-Zaffran et al., 2001). We found that Lar is expressed in R7 photoreceptors during early pupal stages, and single-cell RNA sequencing also revealed that *Lar* is expressed in R7 photoreceptors (Davis et al., 2020). We did not detect Sns reporter expression in R7 photoreceptors, and single-cell RNA sequencing also showed that R7 photoreceptors do not express *sns* (Davis et al., 2020). Hence, Lar and Sns are likely to bind to each other in *trans* to regulate R7 targeting. Sns-expressing neurons arborize in layer M5, but not in M6. Perhaps Sns expression in neurons projecting to M5 provides guidance cues that the R7 terminals employ to remain in the M6 layer.

Lar and Sns are also expressed in several other neuronal types in the optic lobe. One example of a synaptically connected pair of neurons that express Lar and Sns are the L1 and C2 neurons. L1 lamina neurons express Lar>GFP and C2 neurons express Sns>GFP (Fig. 7, Supp. Figs. 6,7). Although *Lar* RNA is expressed at some level in most neurons, RNA sequencing studies have shown that L1 lamina neurons express Lar at particularly high levels (Davis et al., 2020; Tan et al., 2015). L1 neurons send axonal projections to M1 and M5 layers of the medulla (Takemura et al., 2013; Takemura et al., 2015). C2 neurons have dendritic projections in the same layers in the medulla, and they are bidirectionally connected to L1. C2 receives more synapses from L1 than from any other neuron, and also makes many synapses onto L1 (Takemura et al., 2013; Takemura et al., 2015). It is attractive to speculate that Lar-Sns interactions might be important for formation of these synaptic connections. Single-cell RNA sequencing data shows that *kirre* and *rst* are also expressed in L1 neurons (Kurmangaliyev et al., 2020). It is possible that Sns acts by *trans* binding to both Lar and Kirre/Rst in L1 neurons.

### Evolutionary conservation of the Lar-Sns interaction

Sns is a highly conserved CAM that has orthologs in *C. elegans* and mammals (Shen, 2004). The *C. elegans* ortholog of Sns is SYG-2, which has been shown to be required for formation of synapses by the HSNL neuron (Chia et al., 2014; Shen et al., 2004). SYG-2 is expressed on guidepost epithelial cells, and its interactions with SYG-1 on the HSNL neuron initiate the process of synapse formation (Shen et al., 2004). In *SYG-2* and *SYG-1* mutants, components of the presynaptic active zone assembly do not localize properly. This phenotype is similar to that seen in *SYD-2* mutants. SYD-2 is the *C. elegans* ortholog of Liprin-α, which binds to Lar’s cytoplasmic domain. SYG-1/SYG-2 interactions recruit SYD-2 to the site of active zone assembly (Patel et al., 2006) and SYD-2 is mislocalized in mutants for PTP-3, the *C. elegans* Lar ortholog (Ackley et al., 2005). These observations suggest a model of synapse formation in which SYG-1, SYG-2, PTP-3 and SYD-2 all act together at the site of synapse assembly. Replacing worm SYG-1 and SYG-2 with chimeras containing the first IgSF (D1) domains of *Drosophila* and mammalian orthologs was able to rescue the synaptogenesis defects seen in *SYG-1; SYG-2* double mutants (Ozkan et al., 2014).

The mouse and human mammalian Type IIa RPTP subfamily has three members: PTPRF (Lar), PTPRD, and PTPRS. The three Type IIa RPTPs have been shown to bind to an overlapping set of postsynaptic binding partners that are listed in the Introduction. Each of these ligands localizes to the postsynaptic membrane, where they form heterophilic *trans* complexes with presynaptic RPTPs to regulate cell-cell adhesion, presynaptic differentiation and excitatory synapse development (Takahashi and Craig, 2013). The three Type IIa RPTPs have different patterns of expression in the developing mouse brain, with higher expression of LAR and PTPRS compared to PTPRD during early development. Both LAR and PTPRD transcripts are highly enriched in the dorsal root ganglia, and all three are also expressed in the developing mouse spinal cord. LAR is expressed at very low levels in the adult mouse brain, while PTPRS and PTPRD show substantial levels of expression in the adult mouse hippocampus, cerebellum and olfactory bulb (Schaapveld et al., 1998).

*NPHS1* mRNA, encoding Nephrin, is expressed in the spinal cord during early embryonic development until E12 (Putaala et al., 2001). In newborn mice, *NPHS1* is expressed in the cerebellum and several glomeruli of the olfactory bulb, while in adult mice *NPHS1* expression is upregulated in the cerebellum, hippocampus and some glomeruli of the olfactory bulb. The expression patterns of Type IIa *Rptp* and *NPHS1* mRNAs in the developing and adult mouse brain are strikingly similar, with both being expressed in the hippocampus, cerebellum and olfactory bulb. In addition to *Nephrin* mRNA, *Neph1/Kirrel1* mRNA (encoding a mammalian ortholog of Kirre/Rst) is also expressed in the developing mouse neocortex and hippocampus (Volker et al., 2012). Interestingly, Nephrin expression was also observed in the corpus callosum in developing mice (Li and He, 2015), and PTPRS-deficient mice show hippocampal dysgenesis and reductions in the thickness of the corpus callosum (Meathrel et al., 2002). These observations point to a possible role of Nephrin in regulating the developmental functions of Type IIa RPTPs in the mammalian brain.

## Experimental Procedures

### Drosophila genetics

Wild-type (WT) flies used were *Canton S* (CS). *Lar* and *sns* mutants have been previously described: *Lar*^*13.2*^, *Lar*^*451*^ and *sns*^*xb3*^ (gift of Dr. Susan Abmayr). The following lines were obtained from the Bloomington Stock Center: *sns^Df^*, C155-GAL4, tubulin-GAL4, *sns^MiMIC^ ^MI03001^*, *Lar^MiMIC02154^*, *sns^EY08142^*, UAS-Lar RNAi (TRiP.HMS02186) and UAS-Lar RNAi (TRiP.HMS00822). Sns RNAi lines were from the Vienna Drosophila Resource Center: UAS-Sns RNAi (KK109442) and UAS-Sns RNAi (GD877). T2A-GAL4 lines were generated as described in (Diao et al., 2015). Briefly, flies carrying the MiMIC insertion were crossed with flies bearing the triplet “Trojan exon” donor. The F_1_ males from this cross carrying both genetic components were crossed to females carrying germline transgenic sources of Cre and ϕC31. The F_2_ males from this cross that had all four genetic components were then crossed to a UAS-2xEGFP reporter line and the resulting progeny were screened for T2A-GAL4 transformants.

### Immunohistochemistry

Live-dissections of embryos were performed as described in (Lee et al., 2013). Briefly, egg-laying chambers were set up with adult flies and grape juice plates and left in dark at room temperature to lay eggs for four hours. Embryos on the grape plate were incubated overnight at 18°C, followed by two hours at 29°C to induce GAL4 expression. Stage 16 embryos were dissected in PBS, followed by incubation with AP_5_ fusion protein supernatants (1X or concentrated) from lepidopteran HiFive cells infected with baculovirus vectors, transfected Schneider 2 (S2) cells, or transfected mammalian Expi293 cells for two hours at room temperature. Embryos were then fixed in 4% paraformaldehyde for 30 minutes, followed by blocking in 5% normal goat serum in 0.05% PBT (1x PBS with 0.05% Triton-X100 and 0.1% BSA). Primary antibody incubation was done overnight at 4°C. Primary antibodies used were: Rabbit anti-AP (1:1000, AbD Serotec), Mouse anti-FasII (1:3, mAb 1D4, DSHB) and Mouse anti-LAR (1:3, mAb 9D8). Following washes in 0.05% PBT, embryos were incubated in Secondary antibodies for two-four hours at room temperature. Secondary antibodies used were: Alexa Fluor 568 conjugated Goat anti-rabbit and Alexa Fluor 488 conjugated Goat anti-mouse (1:500, Molecular Technologies). Samples were washed and mounted in Vectashield (Vector Laboratories).

Larval dissections were performed as described in (Menon et al., 2015). Briefly, wandering third instar larvae were dissected in PBS and fixed in 4% paraformaldehyde for 30 minutes. Samples were washed in 0.25% PBT (1x PBS with 0.25% Triton X-100 and 0.1% BSA) three times and incubated overnight in 0.25% PBT at 4°C. Samples were blocked in 5% normal goat serum (in 0.25% PBT) for 1-2 hours at room temperature, followed by incubation with primary antibodies overnight at 4°C. Primary antibodies used were: Rabbit anti-GFP (1:500, Molecular Technologies), Rabbit anti-RFP (1:500, Rockland Inc), Mouse anti-Repo (1:10, mAb 8D12, DSHB), Mouse anti-Eve (1:10, mAb 3C10, DSHB), Mouse anti-chaoptin (1:10, mAb 24B10, DSHB), Mouse anti-discs large (1:100, mAb 4F3, DSHB) and Mouse anti-FasII (1:3, mAb 1D4). Following washes in 0.25% PBT, samples were incubated with secondary antibodies overnight at 4°C. Secondary antibodies used were: Alexa Fluor 488 goat anti-rabbit, Alexa Fluor 568 goat anti-rabbit, Alexa Fluor 568 goat anti-mouse, Alexa Fluor 647 goat anti-mouse (1:500, Molecular technologies), Alexa Fluor 488 conjugated goat anti-horseradish peroxidase, Alexa Fluor 568 conjugated goat anti-horseradish peroxidase and Alexa Fluor 647 conjugated goat anti-horseradish peroxidase (1:50, Jackson Immunoresearch). Samples were washed and mounted in Vectashield.

Pupal and adult brain dissections were performed as described in (Menon et al., 2019; Xu et al., 2018). Both pupal and adult brains were dissected in PBS and fixed in 4% paraformaldehyde (in 0.25% PBT) for 30 minutes. Samples were washed overnight at 4°C, followed by blocking in 5% normal goat serum in 0.25% PBT for 1-2 hours at room temperature. Samples were incubated in primary antibodies for two days at 4°C. Primary antibodies used were: Rabbit anti-GFP (1:500, Molecular Technologies), Mouse anti-chaoptin (1:10, mAb 24B10, DSHB), Mouse anti-FasII (1:3, mAb 1D4) and Mouse anti-Trio (1:20, mAb 9.4A, DSHB). Samples were washed in 0.25% PBT and incubated in secondary antibodies for two days at 4°C. Secondary antibodies used were: Alexa Fluor 488 goat anti-rabbit, Alexa Fluor 568 goat anti-mouse, Alexa Fluor 647 goat anti-mouse (1:500, Molecular technologies) and Alexa Fluor 647 conjugated goat anti-horseradish peroxidase (1:50, Jackson Immunoresearch). Following washes, samples were incubated in Vectashield for a minimum of 24 hours before mounting.

### Molecular Biology

AP_5_ and Fc fusion proteins were prepared as described in (Lee et al., 2013) (baculovirus) or in (Ozkan et al., 2013) (S2 cells). S2 cell expression vectors containing the ECDs of Sns and Lar were obtained from the Özkan collection. To make Nephrin, PTPRD, PTPRF and PTPRS AP_5_ and Fc fusion proteins, ECD regions of each were amplified by PCR from full length cDNAs and moved into pCE2 and pCE14 expression vectors using Gateway Cloning. S2 cells were transfected using Effectene transfection reagent as described in (Ozkan et al., 2013). Expression was induced using 100mM CuSO_4_ 24 hours after transfection and supernatants (sups) containing fusion proteins were collected three days after induction. Sups were directly used at 1x concentration for ECIA assays and were concentrated 2-5x using Amicon Ultra-15 centrifugal filter units (30KDa molecular weight cut-off) for use in embryo staining experiments.

### ECIA

Each well of Nunc Maxisorp 96-well plate was incubated with 50 μl of mouse anti-human IgG (Fc-specific) antibody (5 μg/ml in bicarbonate coupling buffer, pH 8.4) overnight at 4°C. Wells were washed in PBST (PBS with 0.05% Tween-20) three times for five minutes each, followed by blocking in 2% BSA (in PBS) for two hours at room temperature. 50 μl Fc fusion proteins were added at 1x concentration for three hours at room temperature, followed by washes and blocking for 30 minutes. 50 μl AP_5_ fusion proteins were added at 1x concentration, pre-clustered with mouse anti-human AP:biotin conjugated antibody (1:500, eBioscience) and incubated overnight at room temperature. Wells were washed in PBST, followed by incubation with Streptavidin: HRP (1:500, 50 μl per well) for 30 minutes. Wells were washed and incubated with 1-Step Ultra TMB HRP substrate (50 μl per well, ThermoFisher) for 30 minutes protected from light. The HRP reaction was stopped by adding 2M phosphoric acid (50 μl per well) and absorbance was measured at 450 nM.

### Confocal imaging and image analysis

All images were captured using a Zeiss LSM710 confocal microscope with either 20X or 40X objectives. NMJs were analyzed using a semi-automated macro in FIJI (Nijhof et al., 2016). 1b and 1s boutons were separately outlined in confocal projections and separate analyses were performed on both kinds of boutons. Dlg immunostaining was used to separate 1b and 1s boutons as 1b boutons stain brightly with Dlg and 1s boutons have very weak Dlg signals. Brp punctae were also counted using the FIJI macro. For MB medial lobe, β and β’ lobe phenotypes, every confocal slice was individually analyzed for FasII-positive axons crossing the midline. For dorsal, α and α’ lobe phenotypes, confocal projections of the entire MB were analyzed for presence or absence of lobes. For R7 photoreceptor targeting phenotype, R7 terminals in M6 layer were counted in at least ten slices per optic lobe with each slice being 5 μm apart. The number of R7 terminals in M6 layer were divided by total number of R7 axons seen in M3 layer and above. Images were analyzed and processed using FIJI software.

### Statistical Analysis

Data were analyzed using Graphpad Prism. For all experiments with the exception of MB phenotypes, statistical analyses were performed using One-Way ANOVA followed by Tukey’s post-hoc correction. MB phenotypes were analyzed using Fisher’s exact test and each genotype was compared to every other genotype from the same experiment. Box and Whisker plots show 10-90 percentile whisker span. For embryo binding experiments, sample size was 8-10 embryos per genotype. For NMJ phenotypes, sample size was 30-60 NMJs per genotype. For larval mushroom body phenotypes, sample size was 12-20 animals per genotype. For adult mushroom body phenotypes, sample size was 20-30 animals per genotype. For optic lobe phenotype, sample size was 10-12 optic lobes per genotype. Each experiment was repeated at least three times.

## Acknowledgments

We thank Michael Anaya for discussions about *in vitro* binding assays, and Kaushiki Menon and Shuwa Xu for general discussions. We thank Susan Abmayr for *sns* lines. This work was supported by a grant from the NIH to K.Z., R37 NS28182.

## Supplemental Figure Legends

**Supplemental Figure 1:**
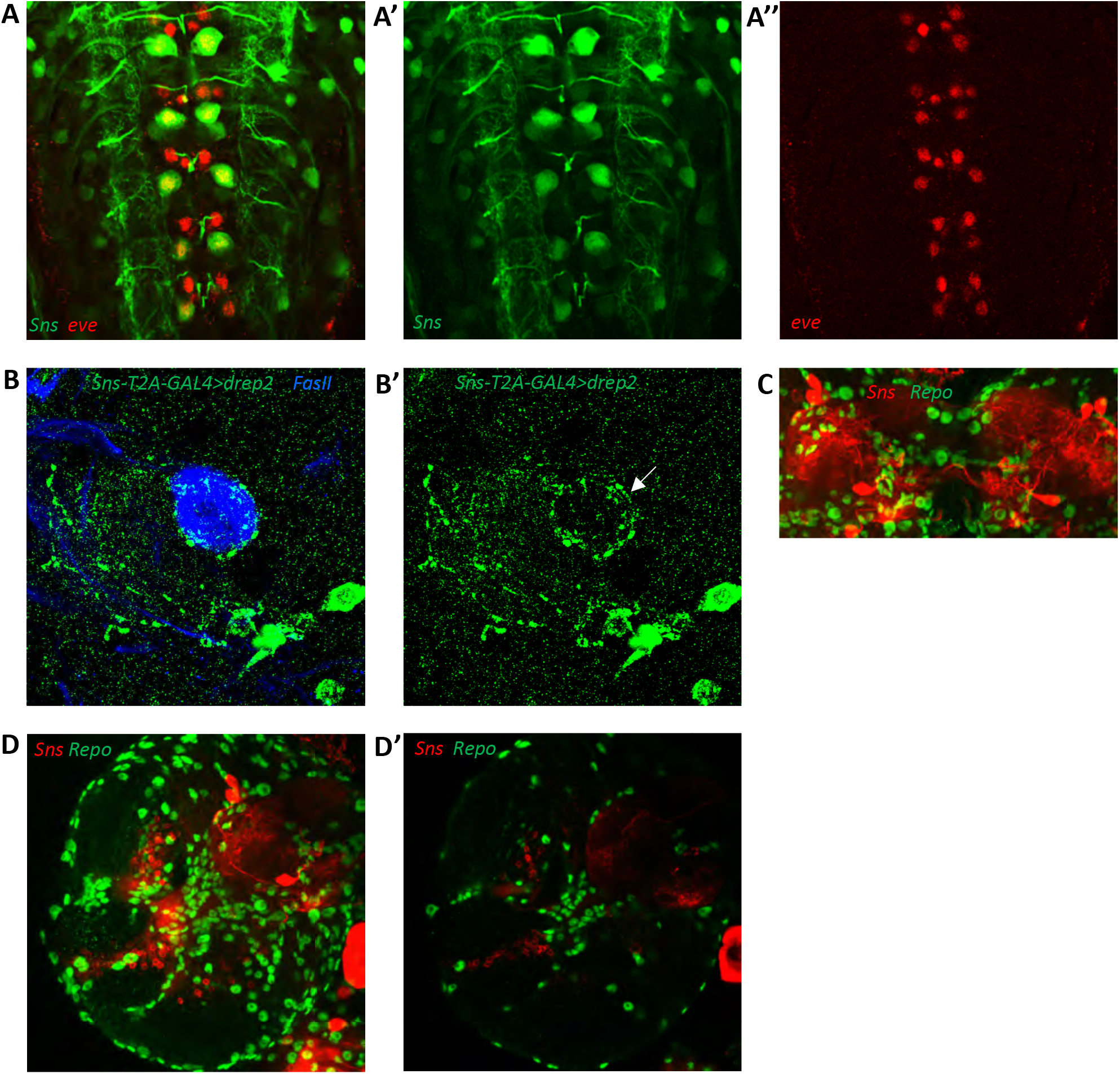
Lar and Sns reporter expression in the larval VNC and brain. (A-A’’) Sns expression visualized by EGFP reporter expression driven by Sns^MI03001-T2A-GAL4^ (Sns>GFP; green), co-stained with Even-skipped antibody (Eve) in the larval VNC. Projection of 4 optical slices is shown. Sns is expressed in Eve-positive RP2/MNISN-1s neurons. (B, B’) Dendritic projections of Sns-expressing neurons seen by driving Drep2 (dendrite-specific marker) with Sns-GAL4 (green), co-stained with FasII antibody to label the mushroom body neuropil (blue). Dendritic projections of Sns-expressing neurons are seen enveloping the dorsal lobe of the mushroom body (B’, arrow). (C-D’) Sns is not expressed in glial cells. (C) Projection of confocal slices through the mushroom body showing no Sns co-expression in Repo-positive glial cells (green). Sns is visualized by driving mCD8-RFP reporter expression using Sns-GAL4 (red). (D, D’) Projection of confocal slices through the entire larval brain, showing no Sns expression in glial cells. Single slice is shown in D’.

**Supplemental Figure 2:**
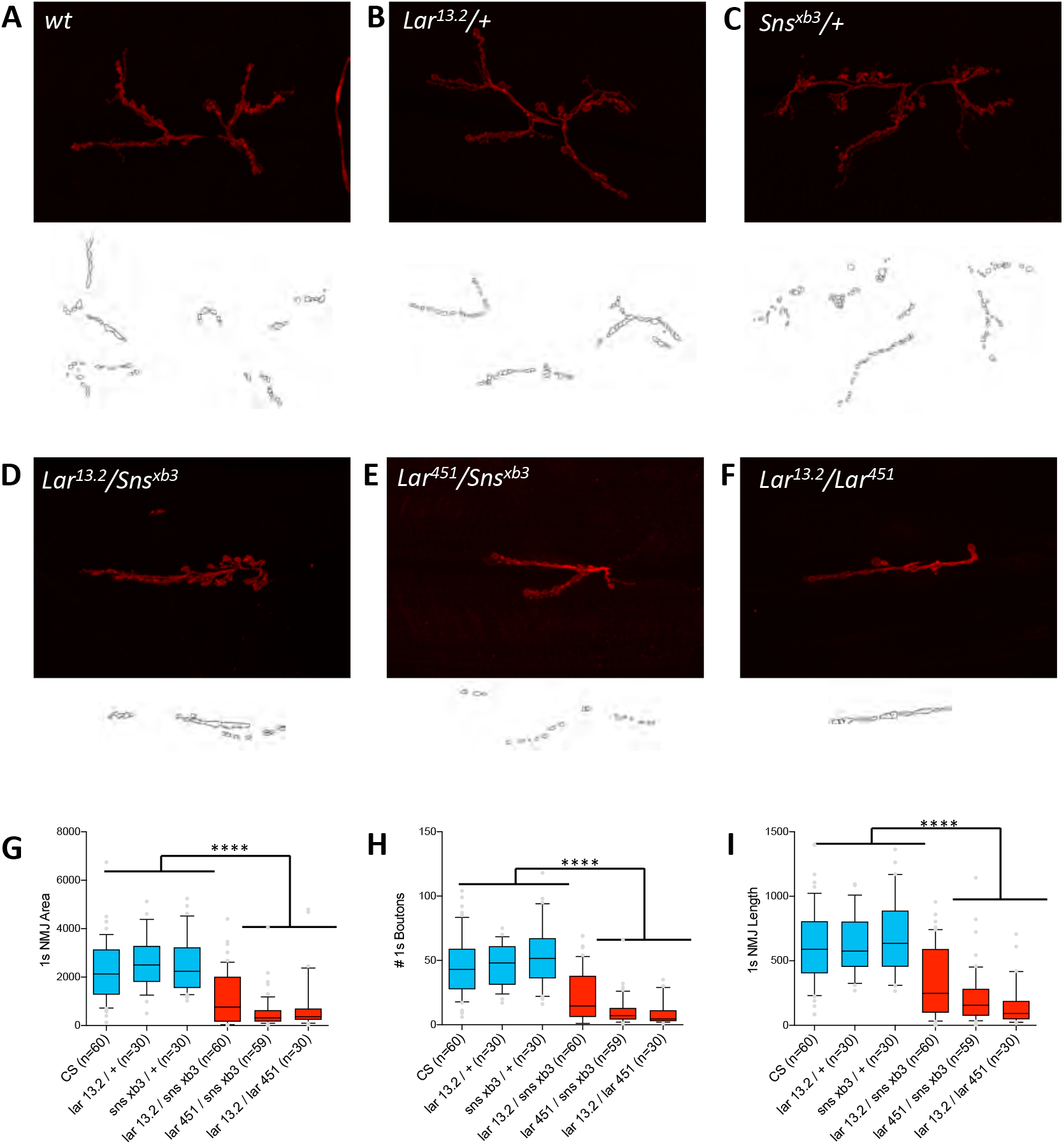
1s NMJ abnormalities in Lar/Sns transhets. NMJs were analyzed using a published FIJI Macro (Nijhof et al., 2016) that uses HRP to outline boutons. NMJ area, length and number of boutons were quantified specifically for 1s boutons. (A-F) Representative images of the NMJ on muscles 7/6 from control (A-C) and transhet (D-F) animals. NMJs are labeled with HRP (red). NMJ outlines showing 1s boutons and branch architecture as outputs from the Macro are under each NMJ image. (G-I) Quantification of 1s NMJ parameters showing reduced NMJ size and boutons in *Lar/Sns* transhets and *Lar* mutants (red) compared to het controls (blue). Data is average from segments A2-A4 from minimum 30 NMJs per genotype. All datasets were analyzed using One-Way ANOVA followed by Tukey’s post-hoc correction. **** p<0.0001.

**Supplemental Figure 3:**
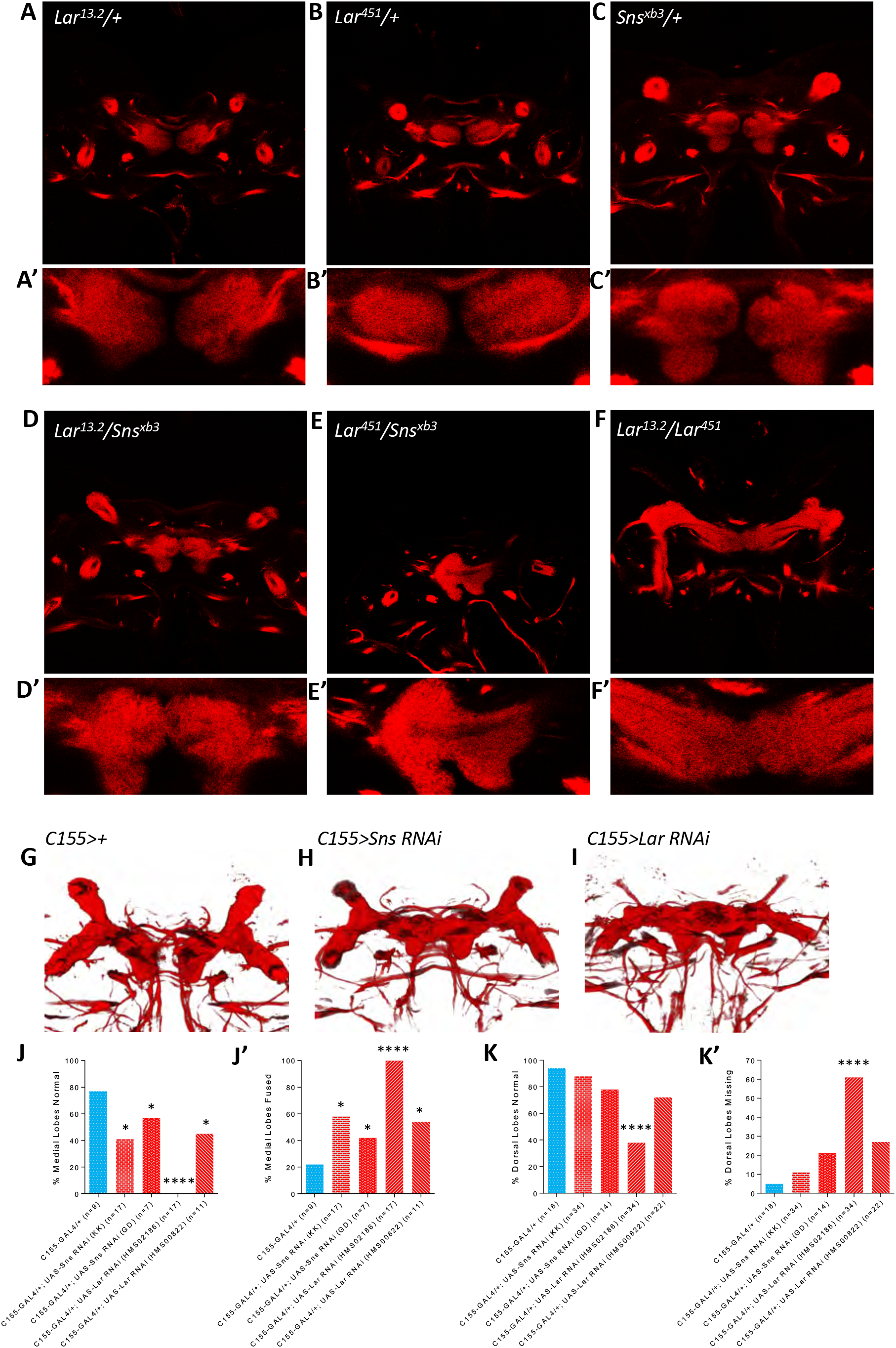
Medial lobe fusion in the larval mushroom body in *Lar/sns* transhets and upon RNAi mediated Lar and Sns knockdown. (A-F’) Single optical slices through the medial lobes of control (A-C’) and genetic transhet (D-F’) animals showing normal unfused medial lobes in controls (close-up of medial lobes in A’-C’), and fused medial lobes in transhets (D’-F’). Anti-FasII (red) is used to label the mushroom body neuropil. (G-I) 3D rendering of confocal stacks of FasII stained larval brains from RNAi knockdown experiments. (J-K’) Quantification of medial lobe fusion and dorsal lobe branching defects upon RNAi-mediated knockdown of Lar and Sns. GAL4 only control in blue and *Lar* or *sns* RNAi genotypes in red. Data were analyzed using Fisher’s Exact Test and each genotype was compared to every other genotype. **** p<0.0001; * p<0.05.

**Supplemental Figure 4:**
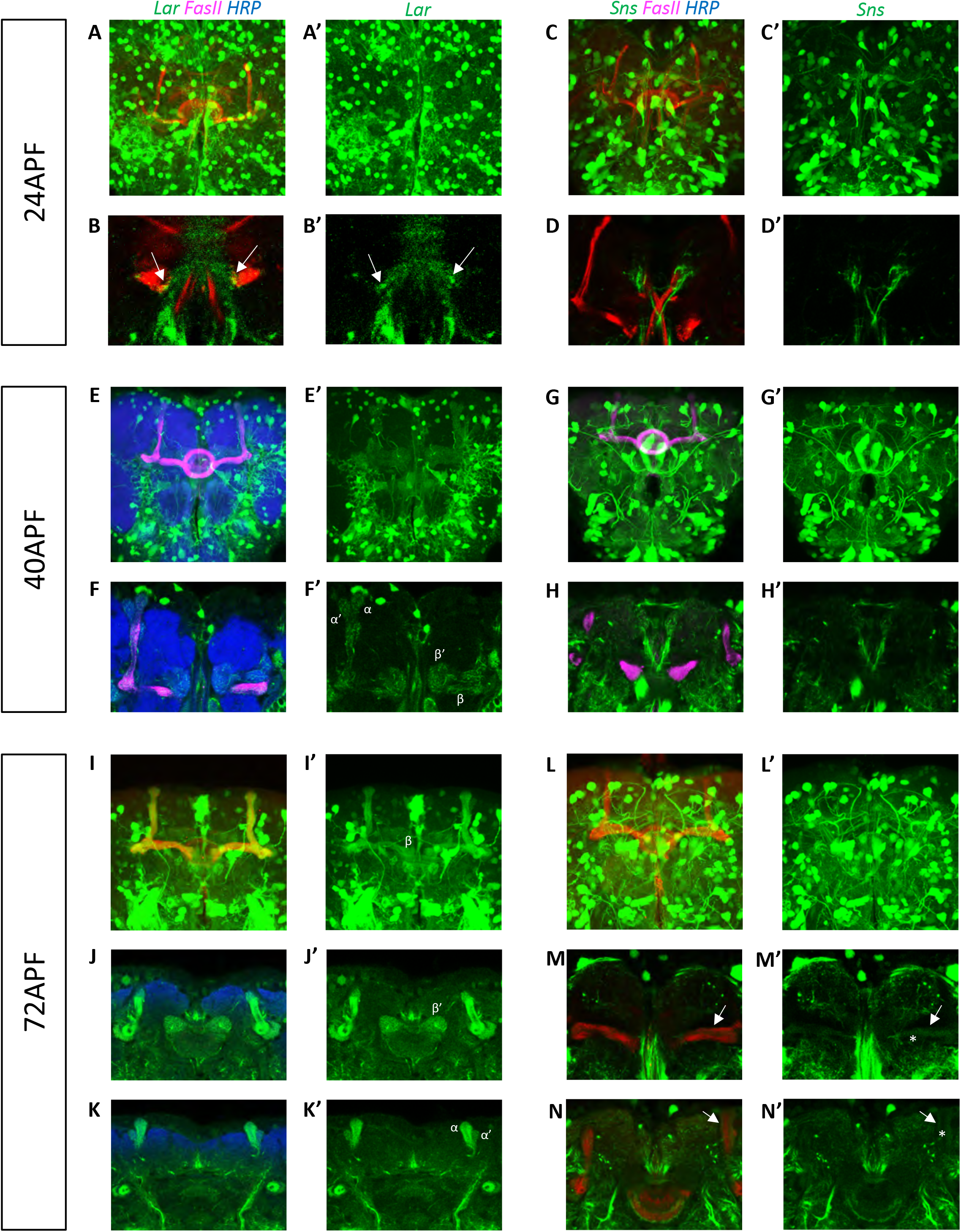
Lar and Sns expression in the developing pupal mushroom body. Confocal projections and single optical slices showing Lar and Sns expression in the 24APF (A-D’), 40APF (E-H’) and 72APF (I-N’) pupal mushroom body, co-stained with FasII antibody (red or magenta). Projections of the entire mushroom body are shown in A, A’, C, C’, E, E’, G, G’, I, I’, L and L’. The rest are single optical slices. Lar expression is seen in the growth cones of developing b lobe axons (B, B’, arrows). No Sns expression is seen in the 24APF mushroom body (C-D’). (E-F’) Lar expression is seen in a, a’, b and b’ lobes in the 40APF mushroom body. (G-H’) No Sns expression is seen in the 40APF mushroom body. (I-K’) Strong Lar expression is seen in all lobes of the 72APF mushroom body. (L-N’) Weak Sns expression is seen in a and b lobes of the 72APF mushroom body (M-N’, arrows). Expression is seen only in older axons in the a and b lobes with hollow cores in the center of the lobes (M’, N’, asterisks).

**Supplemental Figure 5:**
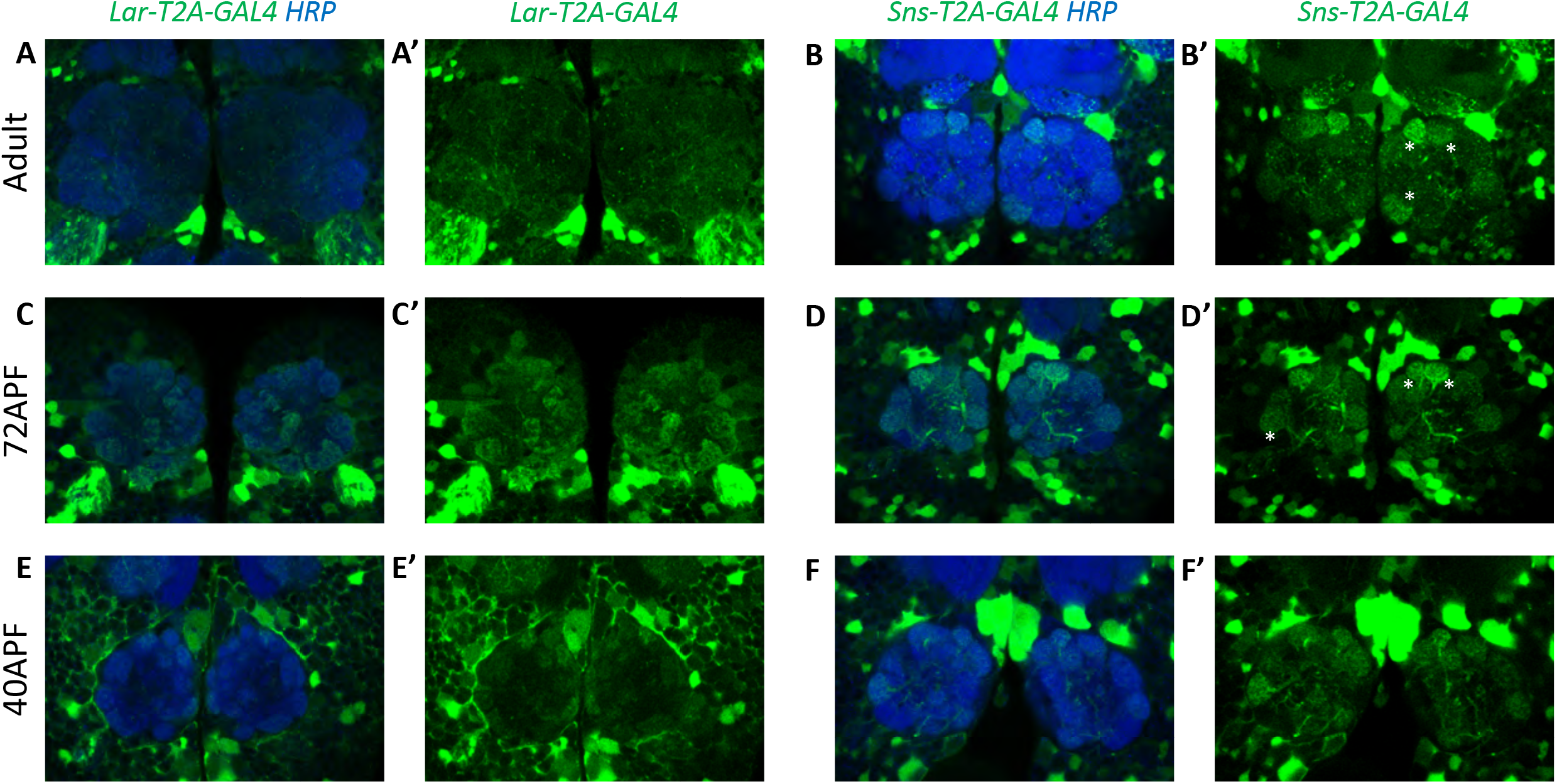
Lar and Sns expression in the adult and developing pupal antennal lobes. Single optical slices showing either Lar>GFP or Sns>GFP (green) expression in the antennal lobes (AL), co-stained with anti-HRP (blue) to label the AL neuropil. (A, A’) Weak Lar>GFP expression seen in several glomeruli of the adult AL. (B, B’) Strong Sns>GFP expression seen in 3 glomeruli of the adult AL (asterisks). (C, C’) Strong Lar>GFP expression seen in several glomeruli in the 72 hr. APF AL. (D, D’) Sns>GFP is expressed at high levels in some glomeruli in the 72 hr. APF AL (asterisks). (E, E’) Weak Lar>GFP expression is seen in several glomeruli in the 40 hr. APF AL. (F, F’) Sns>GFP is strongly expressed in several glomeruli in the 40APF AL.

**Supplemental Figure 6:**
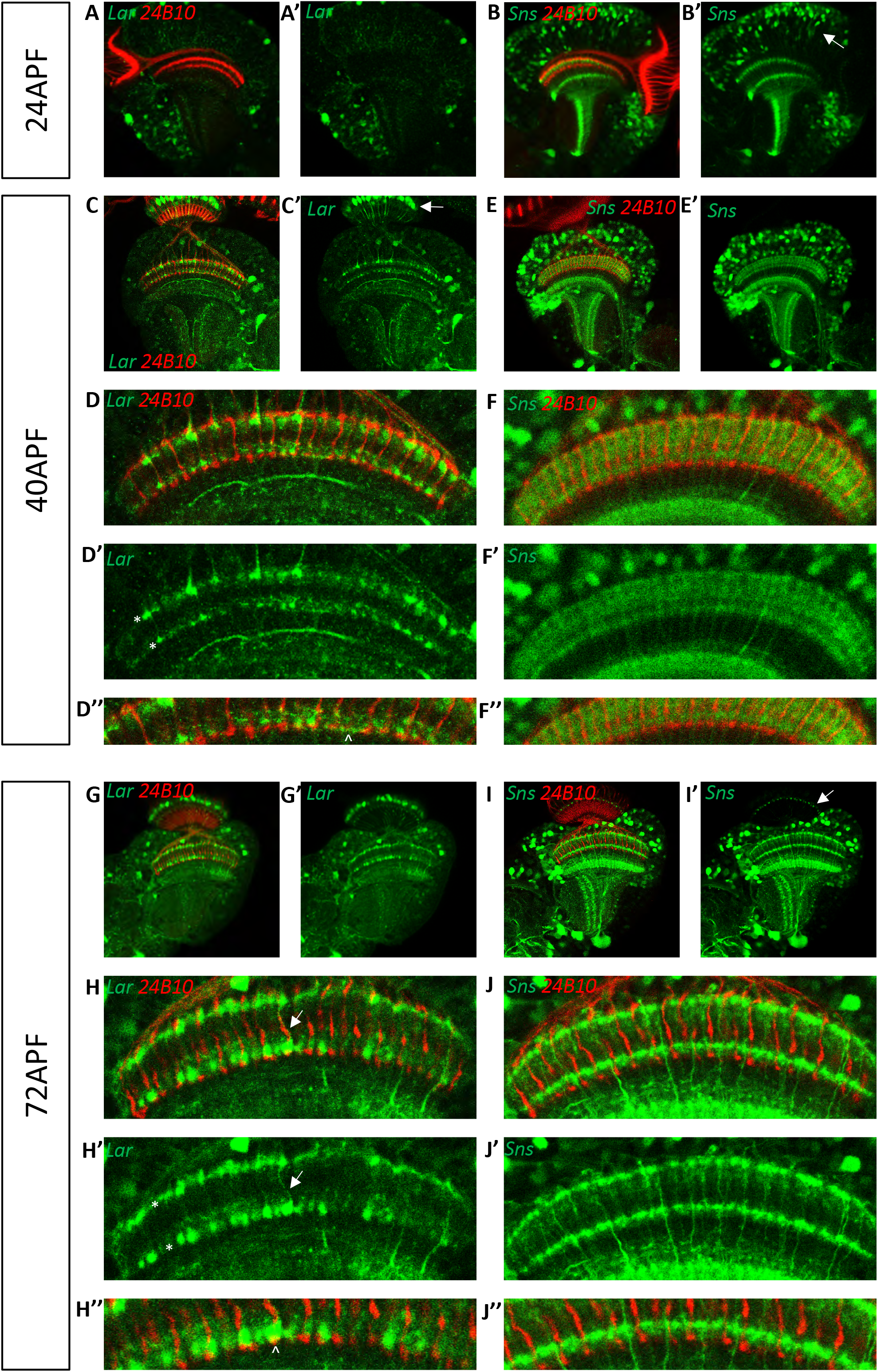
Lar and Sns expression in the developing pupal optic lobes. Single optical slices showing Lar and Sns (green) expression in 24APF, 40APF and 72APF OL, co-stained with anti-Chaoptin (24B10 mAb, red). (A, A’) Weak Lar>GFP expression is seen in the medulla neuropil at 24 hr. APF. (B, B’) At this timepoint, Sns>GFP is expressed at high levels in neuronal cell bodies in the medullary cortex (B’, arrow) and in specific layers in the medulla and lobula. (C, C’) Strong Lar>GFP expression is seen in L1 lamina neuron cell bodies (C’, arrow), which arborize in layers M1 and M5 of the medulla in the 40 hr. APF OL. (D-D’’) Close-up of the distal medulla showing L1 lamina neuron arbors in M1 and M5 layers of the medulla (D’, asterisks). Faint Lar expression is seen in the M6 layer of the medulla (D’’, arrowhead). (E, E’) Sns>GFP expression increases at 40 hr. APF, with many more neurons expressing Sns in the medullary cortex. Sns>GFP expression can be seen in several layers in the distal as well as the proximal medulla. (F-F’’) Close-up of the distal medulla showing Sns>GFP expression in layers M1 through M5 of the medulla. Sns is not expressed in R7 photoreceptors. (G-H’’) Strong Lar>GFP expression seen in L1 cell bodies and layers M1 and M5 of the medulla (H’, asterisks). Strong Lar expression is seen in close proximity to R7 terminals (H’’, arrowhead), and there is labeling of some R7 axons (H’, arrow). (I-J’’) Sns>GFP is expressed at very high levels in the 72 hr. APF optic lobe. Specific Sns expression is seen in M1, M5 and M10 layers of the medulla and a few layers in the lobula (I, I’). Sns expression is also seen in the lamina. Note the dots at the top of the lamina (arrow in I’), which match the morphologies of C2 endings. C2 arborizes in layers M1, M5, and M10 of the medulla (J-J’’) Close-up of the distal medulla showing Sns expression in M1 and M5 layers. Sns is not expressed in R7 photoreceptors.

**Supplemental Figure 7:**
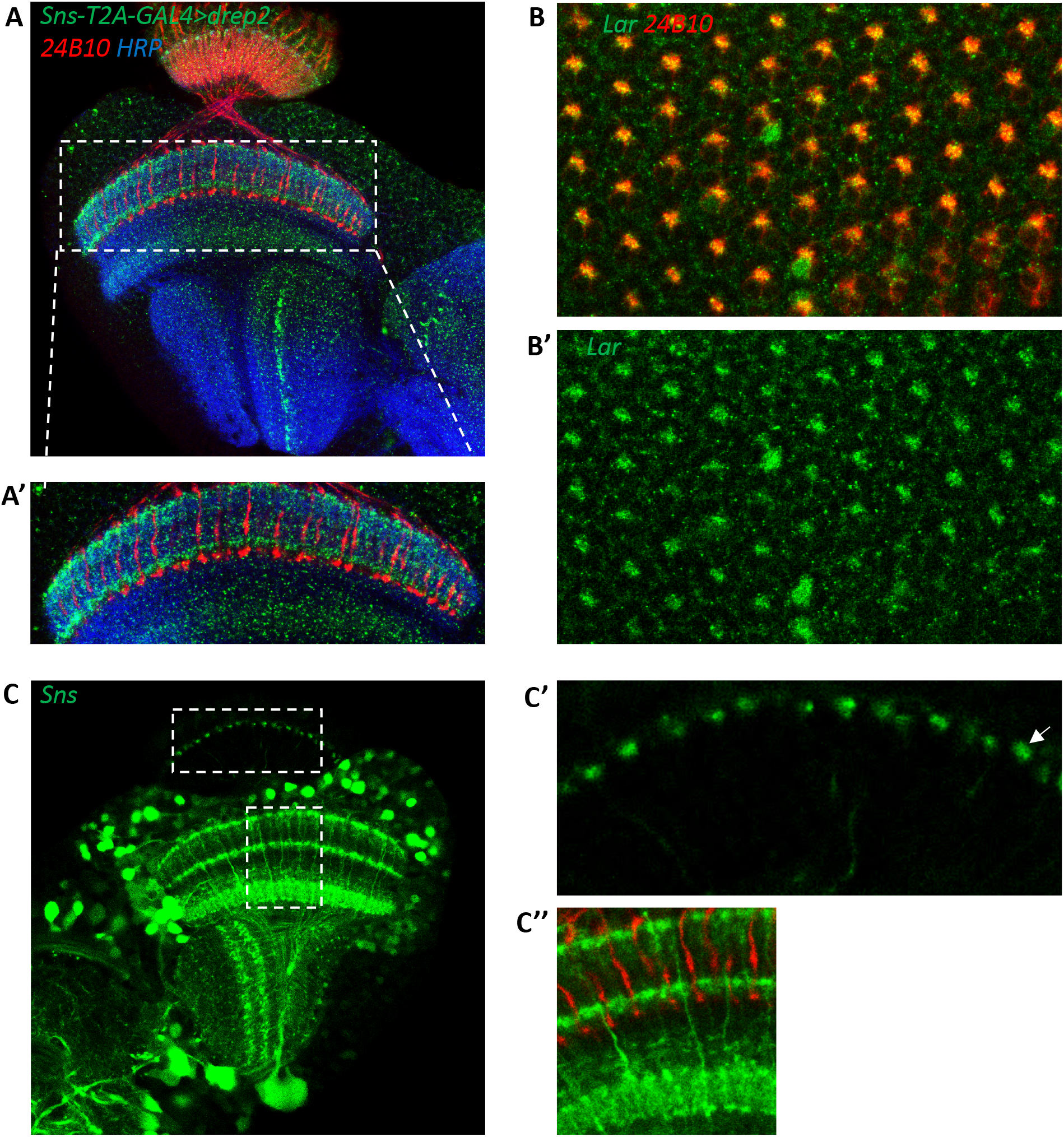
Dendritic projections of Sns neurons in the adult optic lobe and Lar>GFP expression in the 24APF retina. (A’ A’) Single optical slice showing dendritic projections of Sns-expressing neurons visualized by Sns-GAL4>Drep2 (dendrite specific marker, green), co-stained with 24B10 (red) and anti-HRP (blue). Specific dendritic projections can be seen in M1 and M5 layers of the medulla (A’) and in the lamina neuropil. C2 is known to receive synapses in M1, M5, and the lamina. A single layer is seen in the lobula. (B, B’) Lar expression is seen in a top-down view of the 24 hr. APF retina, co-stained with 24B10, (red). R7 and R8 photoreceptor cell bodies lie in the center of each ommatidium. Lar expression is seen in R7 and/or R8 cell bodies in the center of the ommatidia as well as in at least one outer photoreceptor (R1-R6). (C-C’’) Single optical slice showing Sns reporter expression in the OL at 72 hr. APF. Entire optic lobe is shown in (C). (C’) shows close-up of the lamina. Note the dot-like endings of Sns-expressing neurons in the lamina neuropil. These endings most closely resemble those of C2 neurons (Tuthill et al., 2013). (C’’) Sns reporter expression seen in M1, M5 and M10 layers of the medulla, resembling C2 arborization pattern in the medulla (Tuthill et al., 2013).

## Notes

### Competing Interest Statement

The authors have declared no competing interest.

## References

Ackley, B.D., Harrington, R.J., Hudson, M.L., Williams, L., Kenyon, C.J., Chisholm, A.D., and Jin, Y. (2005). The two isoforms of the Caenorhabditis elegans leukocyte-common antigen related receptor tyrosine phosphatase PTP-3 function independently in axon guidance and synapse formation. J Neurosci 25, 7517–7528.

Andlauer, T.F., Scholz-Kornehl, S., Tian, R., Kirchner, M., Babikir, H.A., Depner, H., Loll, B., Quentin, C., Gupta, V.K., Holt, M.G., et al. (2014). Drep-2 is a novel synaptic protein important for learning and memory. Elife 3.

Aricescu, A.R., McKinnell, I.W., Halfter, W., and Stoker, A.W. (2002). Heparan sulfate proteoglycans are ligands for receptor protein tyrosine phosphatase sigma. Mol Cell Biol 22, 1881–1892.

Bao, S., Fischbach, K.F., Corbin, V., and Cagan, R.L. (2010). Preferential adhesion maintains separation of ommatidia in the Drosophila eye. Dev Biol 344, 948–956.

Bour, B.A., Chakravarti, M., West, J.M., and Abmayr, S.M. (2000). Drosophila SNS, a member of the immunoglobulin superfamily that is essential for myoblast fusion. Genes Dev 14, 1498–1511.

Bulgakova, N.A., Klapholz, B., and Brown, N.H. (2012). Cell adhesion in Drosophila: versatility of cadherin and integrin complexes during development. Curr Opin Cell Biol 24, 702–712.

Chia, P.H., Chen, B., Li, P., Rosen, M.K., and Shen, K. (2014). Local F-actin network links synapse formation and axon branching. Cell 156, 208–220.

Choi, Y., Nam, J., Whitcomb, D.J., Song, Y.S., Kim, D., Jeon, S., Um, J.W., Lee, S.G., Woo, J., Kwon, S.K., et al. (2016). SALM5 trans-synaptically interacts with LAR-RPTPs in a splicing-dependent manner to regulate synapse development. Sci Rep 6, 26676.

Clandinin, T.R., Lee, C.H., Herman, T., Lee, R.C., Yang, A.Y., Ovasapyan, S., and Zipursky, S.L. (2001). Drosophila LAR regulates R1-R6 and R7 target specificity in the visual system. Neuron 32, 237–248.

Coles, C.H., Jones, E.Y., and Aricescu, A.R. (2015). Extracellular regulation of type IIa receptor protein tyrosine phosphatases: mechanistic insights from structural analyses. Semin Cell Dev Biol 37, 98–107.

Davis, F.P., Nern, A., Picard, S., Reiser, M.B., Rubin, G.M., Eddy, S.R., and Henry, G.L. (2020). A genetic, genomic, and computational resource for exploring neural circuit function. Elife 9.

Diao, F., Ironfield, H., Luan, H., Diao, F., Shropshire, W.C., Ewer, J., Marr, E., Potter, C.J., Landgraf, M., and White, B.H. (2015). Plug-and-play genetic access to drosophila cell types using exchangeable exon cassettes. Cell Rep 10, 1410–1421.

Fischbach, K.F., Linneweber, G.A., Andlauer, T.F., Hertenstein, A., Bonengel, B., and Chaudhary, K. (2009). The irre cell recognition module (IRM) proteins. J Neurogenet 23, 48–67.

Fox, A.N., and Zinn, K. (2005). The heparan sulfate proteoglycan syndecan is an in vivo ligand for the Drosophila LAR receptor tyrosine phosphatase. Curr Biol 15, 1701–1711.

Galletta, B.J., Chakravarti, M., Banerjee, R., and Abmayr, S.M. (2004). SNS: Adhesive properties, localization requirements and ectodomain dependence in S2 cells and embryonic myoblasts. Mech Dev 121, 1455–1468.

Hakeda-Suzuki, S., Takechi, H., Kawamura, H., and Suzuki, T. (2017). Two receptor tyrosine phosphatases dictate the depth of axonal stabilizing layer in the visual system. Elife 6.

Hattori, D., Demir, E., Kim, H.W., Viragh, E., Zipursky, S.L., and Dickson, B.J. (2007). Dscam diversity is essential for neuronal wiring and self-recognition. Nature 449, 223–227.

Hattori, D., Millard, S.S., Wojtowicz, W.M., and Zipursky, S.L. (2008). Dscam-mediated cell recognition regulates neural circuit formation. Annu Rev Cell Dev Biol 24, 597–620.

Hofmeyer, K., and Treisman, J.E. (2009). The receptor protein tyrosine phosphatase LAR promotes R7 photoreceptor axon targeting by a phosphatase-independent signaling mechanism. Proc Natl Acad Sci U S A 106, 19399–19404.

Horn, K.E., Xu, B., Gobert, D., Hamam, B.N., Thompson, K.M., Wu, C.L., Bouchard, J.F., Uetani, N., Racine, R.J., Tremblay, M.L., et al. (2012). Receptor protein tyrosine phosphatase sigma regulates synapse structure, function and plasticity. J Neurochem 122, 147–161.

Jeon, M., Nguyen, H., Bahri, S., and Zinn, K. (2008). Redundancy and compensation in axon guidance: genetic analysis of the Drosophila Ptp10D/Ptp4E receptor tyrosine phosphatase subfamily. Neural Dev 3, 3.

Jeon, M., and Zinn, K. (2015). R3 receptor tyrosine phosphatases: conserved regulators of receptor tyrosine kinase signaling and tubular organ development. Semin Cell Dev Biol 37, 119–126.

Johnson, K.G., Tenney, A.P., Ghose, A., Duckworth, A.M., Higashi, M.E., Parfitt, K., Marcu, O., Heslip, T.R., Marsh, J.L., Schwarz, T.L., et al. (2006). The HSPGs Syndecan and Dallylike bind the receptor phosphatase LAR and exert distinct effects on synaptic development. Neuron 49, 517–531.

Johnson, K.G., and Van Vactor, D. (2003). Receptor protein tyrosine phosphatases in nervous system development. Physiol Rev 83, 1–24.

Kaufmann, N., DeProto, J., Ranjan, R., Wan, H., and Van Vactor, D. (2002). Drosophila liprin-alpha and the receptor phosphatase Dlar control synapse morphogenesis. Neuron 34, 27–38.

Koch, N., Kobler, O., Thomas, U., Qualmann, B., and Kessels, M.M. (2014). Terminal axonal arborization and synaptic bouton formation critically rely on abp1 and the arp2/3 complex. PLoS One 9, e97692.

Kocherlakota, K.S., Wu, J.M., McDermott, J., and Abmayr, S.M. (2008). Analysis of the cell adhesion molecule sticks-and-stones reveals multiple redundant functional domains, protein-interaction motifs and phosphorylated tyrosines that direct myoblast fusion in Drosophila melanogaster. Genetics 178, 1371–1383.

Krueger, N.X., Van Vactor, D., Wan, H.I., Gelbart, W.M., Goodman, C.S., and Saito, H. (1996). The transmembrane tyrosine phosphatase DLAR controls motor axon guidance in Drosophila. Cell 84, 611–622.

Kurmangaliyev, Y.Z., Yoo, J., Valdes-Aleman, J., Sanfilippo, P., and Zipursky, S.L. (2020). Transcriptional Programs of Circuit Assembly in the Drosophila Visual System. Neuron.

Kurusu, M., Cording, A., Taniguchi, M., Menon, K., Suzuki, E., and Zinn, K. (2008). A screen of cell-surface molecules identifies leucine-rich repeat proteins as key mediators of synaptic target selection. Neuron 59, 972–985.

Kurusu, M., and Zinn, K. (2008). Receptor tyrosine phosphatases regulate birth order-dependent axonal fasciculation and midline repulsion during development of the Drosophila mushroom body. Mol Cell Neurosci 38, 53–65.

Kwon, S.K., Woo, J., Kim, S.Y., Kim, H., and Kim, E. (2010). Trans-synaptic adhesions between netrin-G ligand-3 (NGL-3) and receptor tyrosine phosphatases LAR, protein-tyrosine phosphatase delta (PTPdelta), and PTPsigma via specific domains regulate excitatory synapse formation. J Biol Chem 285, 13966–13978.

Lee, H.K., Cording, A., Vielmetter, J., and Zinn, K. (2013). Interactions between a receptor tyrosine phosphatase and a cell surface ligand regulate axon guidance and glial-neuronal communication. Neuron 78, 813–826.

Li, J., Han, S., Li, H., Udeshi, N.D., Svinkina, T., Mani, D.R., Xu, C., Guajardo, R., Xie, Q., Li, T., et al. (2020). Cell-Surface Proteomic Profiling in the Fly Brain Uncovers Wiring Regulators. Cell 180, 373–386 e315.

Li, X., and He, J.C. (2015). An update: the role of Nephrin inside and outside the kidney. Sci China Life Sci 58, 649–657.

Li, Y., Zhang, P., Choi, T.Y., Park, S.K., Park, H., Lee, E.J., Lee, D., Roh, J.D., Mah, W., Kim, R., et al. (2015). Splicing-Dependent Trans-synaptic SALM3-LAR-RPTP Interactions Regulate Excitatory Synapse Development and Locomotion. Cell Rep 12, 1618–1630.

Liu, G., Kaw, B., Kurfis, J., Rahmanuddin, S., Kanwar, Y.S., and Chugh, S.S. (2003). Neph1 and nephrin interaction in the slit diaphragm is an important determinant of glomerular permeability. J Clin Invest 112, 209–221.

Maurel-Zaffran, C., Suzuki, T., Gahmon, G., Treisman, J.E., and Dickson, B.J. (2001). Cell-autonomous and -nonautonomous functions of LAR in R7 photoreceptor axon targeting. Neuron 32, 225–235.

Meathrel, K., Adamek, T., Batt, J., Rotin, D., and Doering, L.C. (2002). Protein tyrosine phosphatase sigma-deficient mice show aberrant cytoarchitecture and structural abnormalities in the central nervous system. J Neurosci Res 70, 24–35.

Menon, K.P., Carrillo, R.A., and Zinn, K. (2015). The translational regulator Cup controls NMJ presynaptic terminal morphology. Mol Cell Neurosci 67, 126–136.

Menon, K.P., Kulkarni, V., Takemura, S.Y., Anaya, M., and Zinn, K. (2019). Interactions between Dpr11 and DIP-gamma control selection of amacrine neurons in Drosophila color vision circuits. Elife 8.

Nijhof, B., Castells-Nobau, A., Wolf, L., Scheffer-de Gooyert, J.M., Monedero, I., Torroja, L., Coromina, L., van der Laak, J.A., and Schenck, A. (2016). A New Fiji-Based Algorithm That Systematically Quantifies Nine Synaptic Parameters Provides Insights into Drosophila NMJ Morphometry. PLoS Comput Biol 12, e1004823.

Ozkan, E., Carrillo, R.A., Eastman, C.L., Weiszmann, R., Waghray, D., Johnson, K.G., Zinn, K., Celniker, S.E., and Garcia, K.C. (2013). An extracellular interactome of immunoglobulin and LRR proteins reveals receptor-ligand networks. Cell 154, 228–239.

Ozkan, E., Chia, P.H., Wang, R.R., Goriatcheva, N., Borek, D., Otwinowski, Z., Walz, T., Shen, K., and Garcia, K.C. (2014). Extracellular architecture of the SYG-1/SYG-2 adhesion complex instructs synaptogenesis. Cell 156, 482–494.

Patel, M.R., Lehrman, E.K., Poon, V.Y., Crump, J.G., Zhen, M., Bargmann, C.I., and Shen, K. (2006). Hierarchical assembly of presynaptic components in defined C. elegans synapses. Nat Neurosci 9, 1488–1498.

Prakash, S., Caldwell, J.C., Eberl, D.F., and Clandinin, T.R. (2005). Drosophila N-cadherin mediates an attractive interaction between photoreceptor axons and their targets. Nat Neurosci 8, 443–450.

Putaala, H., Soininen, R., Kilpelainen, P., Wartiovaara, J., and Tryggvason, K. (2001). The murine nephrin gene is specifically expressed in kidney, brain and pancreas: inactivation of the gene leads to massive proteinuria and neonatal death. Hum Mol Genet 10, 1–8.

Ruotsalainen, V., Ljungberg, P., Wartiovaara, J., Lenkkeri, U., Kestila, M., Jalanko, H., Holmberg, C., and Tryggvason, K. (1999). Nephrin is specifically located at the slit diaphragm of glomerular podocytes. Proc Natl Acad Sci U S A 96, 7962–7967.

Schaapveld, R.Q., Schepens, J.T., Bachner, D., Attema, J., Wieringa, B., Jap, P.H., and Hendriks, W.J. (1998). Developmental expression of the cell adhesion molecule-like protein tyrosine phosphatases LAR, RPTPdelta and RPTPsigma in the mouse. Mech Dev 77, 59–62.

Sens, K.L., Zhang, S., Jin, P., Duan, R., Zhang, G., Luo, F., Parachini, L., and Chen, E.H. (2010). An invasive podosome-like structure promotes fusion pore formation during myoblast fusion. J Cell Biol 191, 1013–1027.

Shelton, C., Kocherlakota, K.S., Zhuang, S., and Abmayr, S.M. (2009). The immunoglobulin superfamily member Hbs functions redundantly with Sns in interactions between founder and fusion-competent myoblasts. Development 136, 1159–1168.

Shen, K. (2004). Molecular mechanisms of target specificity during synapse formation. Curr Opin Neurobiol 14, 83–88.

Shen, K., Fetter, R.D., and Bargmann, C.I. (2004). Synaptic specificity is generated by the synaptic guidepost protein SYG-2 and its receptor, SYG-1. Cell 116, 869–881.

Stepanek, L., Stoker, A.W., Stoeckli, E., and Bixby, J.L. (2005). Receptor tyrosine phosphatases guide vertebrate motor axons during development. J Neurosci 25, 3813–3823.

Stewart, K., Uetani, N., Hendriks, W., Tremblay, M.L., and Bouchard, M. (2013). Inactivation of LAR family phosphatase genes Ptprs and Ptprf causes craniofacial malformations resembling Pierre-Robin sequence. Development 140, 3413–3422.

Sudhof, T.C. (2008). Neuroligins and neurexins link synaptic function to cognitive disease. Nature 455, 903–911.

Sun, Q., Bahri, S., Schmid, A., Chia, W., and Zinn, K. (2000). Receptor tyrosine phosphatases regulate axon guidance across the midline of the Drosophila embryo. Development 127, 801–812.

Takahashi, H., Arstikaitis, P., Prasad, T., Bartlett, T.E., Wang, Y.T., Murphy, T.H., and Craig, A.M. (2011). Postsynaptic TrkC and presynaptic PTPsigma function as a bidirectional excitatory synaptic organizing complex. Neuron 69, 287–303.

Takahashi, H., and Craig, A.M. (2013). Protein tyrosine phosphatases PTPdelta, PTPsigma, and LAR: presynaptic hubs for synapse organization. Trends Neurosci 36, 522–534.

Takahashi, H., Katayama, K., Sohya, K., Miyamoto, H., Prasad, T., Matsumoto, Y., Ota, M., Yasuda, H., Tsumoto, T., Aruga, J., et al. (2012). Selective control of inhibitory synapse development by Slitrk3-PTPdelta trans-synaptic interaction. Nat Neurosci 15, 389–398, S381-382.

Takemura, S.Y., Bharioke, A., Lu, Z., Nern, A., Vitaladevuni, S., Rivlin, P.K., Katz, W.T., Olbris, D.J., Plaza, S.M., Winston, P., et al. (2013). A visual motion detection circuit suggested by Drosophila connectomics. Nature 500, 175–181.

Takemura, S.Y., Xu, C.S., Lu, Z., Rivlin, P.K., Parag, T., Olbris, D.J., Plaza, S., Zhao, T., Katz, W.T., Umayam, L., et al. (2015). Synaptic circuits and their variations within different columns in the visual system of Drosophila. Proc Natl Acad Sci U S A 112, 13711–13716.

Tan, L., Zhang, K.X., Pecot, M.Y., Nagarkar-Jaiswal, S., Lee, P.T., Takemura, S.Y., McEwen, J.M., Nern, A., Xu, S., Tadros, W., et al. (2015). Ig Superfamily Ligand and Receptor Pairs Expressed in Synaptic Partners in Drosophila. Cell 163, 1756–1769.

Tuthill, J.C., Nern, A., Holtz, S.L., Rubin, G.M., and Reiser, M.B. (2013). Contributions of the 12 neuron classes in the fly lamina to motion vision. Neuron 79, 128–140.

Uetani, N., Chagnon, M.J., Kennedy, T.E., Iwakura, Y., and Tremblay, M.L. (2006). Mammalian motoneuron axon targeting requires receptor protein tyrosine phosphatases sigma and delta. J Neurosci 26, 5872–5880.

Uetani, N., Kato, K., Ogura, H., Mizuno, K., Kawano, K., Mikoshiba, K., Yakura, H., Asano, M., and Iwakura, Y. (2000). Impaired learning with enhanced hippocampal long-term potentiation in PTPdelta-deficient mice. EMBO J 19, 2775–2785.

Volker, L.A., Petry, M., Abdelsabour-Khalaf, M., Schweizer, H., Yusuf, F., Busch, T., Schermer, B., Benzing, T., Brand-Saberi, B., Kretz, O., et al. (2012). Comparative analysis of Neph gene expression in mouse and chicken development. Histochem Cell Biol 137, 355–366.

Wojtowicz, W.M., Vielmetter, J., Fernandes, R.A., Siepe, D.H., Eastman, C.L., Chisholm, G.B., Cox, S., Klock, H., Anderson, P.W., Rue, S.M., et al. (2020). A Human IgSF Cell-Surface Interactome Reveals a Complex Network of Protein-Protein Interactions. Cell 182, 1027–1043 e1017.

Woo, J., Kwon, S.K., Choi, S., Kim, S., Lee, J.R., Dunah, A.W., Sheng, M., and Kim, E. (2009). Trans-synaptic adhesion between NGL-3 and LAR regulates the formation of excitatory synapses. Nat Neurosci 12, 428–437.

Xu, S., Xiao, Q., Cosmanescu, F., Sergeeva, A.P., Yoo, J., Lin, Y., Katsamba, P.S., Ahlsen, G., Kaufman, J., Linaval, N.T., et al. (2018). Interactions between the Ig-Superfamily Proteins DIP-alpha and Dpr6/10 Regulate Assembly of Neural Circuits. Neuron 100, 1369–1384 e1366.

Yim, Y.S., Kwon, Y., Nam, J., Yoon, H.I., Lee, K., Kim, D.G., Kim, E., Kim, C.H., and Ko, J. (2013). Slitrks control excitatory and inhibitory synapse formation with LAR receptor protein tyrosine phosphatases. Proc Natl Acad Sci U S A 110, 4057–4062.

Yoshida, T., Shiroshima, T., Lee, S.J., Yasumura, M., Uemura, T., Chen, X., Iwakura, Y., and Mishina, M. (2012). Interleukin-1 receptor accessory protein organizes neuronal synaptogenesis as a cell adhesion molecule. J Neurosci 32, 2588–2600.

Yoshida, T., Yasumura, M., Uemura, T., Lee, S.J., Ra, M., Taguchi, R., Iwakura, Y., and Mishina, M. (2011). IL-1 receptor accessory protein-like 1 associated with mental retardation and autism mediates synapse formation by trans-synaptic interaction with protein tyrosine phosphatase delta. J Neurosci 31, 13485–13499.

Zhuang, S., Shao, H., Guo, F., Trimble, R., Pearce, E., and Abmayr, S.M. (2009). Sns and Kirre, the Drosophila orthologs of Nephrin and Neph1, direct adhesion, fusion and formation of a slit diaphragm-like structure in insect nephrocytes. Development 136, 2335–2344.

